# Iterative Epitope Expansion Enables Structure-Guided *de novo* Design of IgE-Binding Miniproteins Targeting the Fc*ε*RI Interface

**DOI:** 10.64898/2025.12.27.696647

**Authors:** Ido Calman, Hamed Shaykhalishahi, Puwanat Sangkhapreecha, Sharrol Bachas

## Abstract

Designing *de novo* protein binders to multidomain assemblies remains challenging due to their structural complexity, conformational diversity, and broad functional epitopes. Here, we present an iterative epitope-expansion strategy that enables structure-guided design of compact miniproteins targeting human immunoglobulin E (IgE). Direct attempts to design binders against the full Fc*ε*RI-binding interface of IgE produced candidates with poor *in silico* confidence and limited experimental success. Instead, by first targeting a nearby, geometrically tractable seed epitope on the IgE C*_ε_*3 domain, we obtained initial binders that served as anchors for progressive, structure-guided expansion toward the full receptor-binding interface. Iteration through an integrated AI–human–wet-lab design loop yielded a second-round miniprotein with approximately 30-fold improved affinity and substantial structural overlap with the Fc*ε*RI-binding site.

Alanine-scanning mutagenesis confirmed that loss-of-function substitutions mapped precisely to the designed binding interfaces, validating the structural models. Polyspecificity profiling revealed that two of three binders exhibited low off-target binding comparable to antibody benchmarks, whereas one design displayed elevated polyspecificity that was strongly amplified upon Fc fusion. Notably, a previously published high-affinity miniprotein benchmark exhibited similar format-dependent liabilities, underscoring an underappreciated developability risk in the *de novo* miniprotein literature. Competition assays further demonstrated that two binders occupy regions of IgE that overlap with the Fc*ε*RI binding interface, as evidenced by loss of miniprotein binding when Fc*ε*RI is pre-bound to IgE Fc, consistent with the intended design geometry.

Together, these results establish epitope expansion as a generalizable strategy for steering generative protein design toward challenging multidomain epitopes and highlight the necessity of tightly coupled AI–human–experimental workflows for achieving both functional specificity and developability in *de novo* binder discovery.

## Introduction

Recent advances in *de novo* protein design and generative modeling have enabled the creation of compact binders with remarkable structural accuracy [1–15]. Several next-generation approaches—including diffusion-based hallucination frameworks, joint co-folding models, and sequence–structure generative systems—can now design miniproteins and even full antibody variable domains that fold as intended and bind their targets. However, most reported successes involve single-domain proteins or engineered constructs in which the epitope resides on a single polypeptide chain [1, 2, 5, 6, 10, 11, 13, 15]. These methods typically require truncated domains or fragments to fit within model context limits. While highly effective for monomeric or well-behaved domain targets, such strategies do not readily generalize to multimeric assemblies or discontinuous epitopes spanning multiple domains.

This limitation is particularly important because many biologically relevant receptors—including GPCRs, cytokine receptors, and immune co-stimulatory molecules—present composite epitopes shaped by inter-domain geometry. Designing binders that recognize such surfaces requires computational strategies focused on intact multidomain assemblies rather than simplified fragments. In practice, state-of-the-art generative models often struggle in these settings; for example, most models fail to produce viable binders to trimeric TNF-*α* [10, 11, 15], despite partial success reported for the Chai-2 generative model [2], for which epitope specificity has not yet been experimentally validated. These challenges highlight the need for design strategies capable of navigating structurally complex and discontinuous binding surfaces.

To evaluate computational strategies for multidomain epitopes, we selected human immunoglobulin E (IgE) as a case study. IgE is a heavily glycosylated, multidomain antibody isotype that plays a central role in allergic reactions [16, 17]. Through its Fc region, IgE interacts with two immune receptors: Fc*ε*RI and Fc*ε*RII (CD23) [16]. Circulating IgE binds Fc*ε*RI on mast cells and basophils with high affinity, priming these cells for activation [18]. Allergen engagement of the IgE–Fc*ε*RI complex triggers receptor crosslinking, cellular degranulation, and release of histamine and inflammatory mediators [17]. Importantly, the Fc*ε*RI-binding site on IgE spans the C*_ε_*2–C*_ε_*3 interface, forming a composite epitope that cannot be isolated as a single domain [19,20]. This makes the IgE–Fc*ε*RI interface an ideal system for assessing whether modern *de novo* design methods can address challenging multidomain targets. Moreover, the availability of a high-affinity natural receptor provides a rigorous benchmark for structural and functional validation of designed binders [19].

In this work, we employed the Latent-X1 generative model to design binders against the IgE–Fc*ε*RI interface [13]. Latent-X1 has been validated as an efficient engine for generating structurally diverse, high-affinity binders across a range of targets. However, similar to other advanced generative systems, Latent-X1 displayed low confidence metrics when prompted to design directly against the full discontinuous Fc*ε*RI-binding surface. To overcome this limitation, we introduce an epitope-expansion strategy to progressively steer designs toward the desired interface. Rather than generating binders against the entire Fc*ε*RI interface—which produced candidates with poor scoring metrics—we first targeted a nearby, partially overlapping epitope that yielded high-quality latent-space generations. This epitope served as a seed for iterative expansion toward the full Fc*ε*RI site.

Our design strategy relies on an iterative AI–human–wet-lab loop. Latent-X1 generates large pools of candidate binders, which we first screen computationally for epitope alignment, interface geometry, and global structural confidence using a weighted combination of iPTM, predicted aligned error (PAE), and complex RMSD. Human inspection then evaluates surviving models for plausibility, emphasizing interface packing, orientation, and local stereochemistry. Fewer than fifty candidates were selected for round-one validation, and fewer than thirty for the epitope-expansion round. Across both rounds the overall experimental hit rate was approximately 6%, underscoring the importance of strict model triage for multidomain targets.

A key enabler of this workflow was our rapid binder-characterization pipeline, which provides binder expression and affinity assessment within roughly seven days. This fast feedback cycle allowed us to identify initial IgE-binding miniproteins, refine interface hypotheses, and iteratively expand designs toward the Fc*ε*RI epitope. We then carried out deeper experimental validation. Alanine scanning identified the residues essential for binding and mapped them directly onto the designed interface. Polyspecificity profiling by BVP-ELISA clearly distinguished specific binders from variants with off-target liabilities, including one design whose nonspecific reactivity was strongly amplified upon Fc fusion. Finally, competition assays showed that binders engage the Fc*ε*RI interface as intended.

Together, these iterative design and validation cycles enabled successful targeting of a complex, multidomain epitope on full-length IgE Fc. To our knowledge, these results represent the first reported *de novo* miniprotein binders to human IgE and establish an end-to-end framework for designing against structurally challenging epitopes.

## Results

### Selection of the IgE Target Epitope

The human IgE constant region is a multidomain antibody isotype whose Fc region comprises the C*ε*2, C*ε*3, and C*ε*4 domains arranged in a bent, asymmetric orientation [19, 21] (Fig. 1A). The high-affinity receptor Fc*ε*RI recognizes IgE through a composite surface spanning the C*ε*2–C*ε*3 interface [19, 22]. Structural studies show that Fc*ε*RI*α* binds asymmetrically, inserting a concave loop into a shallow pocket at this junction while simultaneously engaging residues on the opposite face of C*ε*3 [19, 22]. These contacts stabilize the bent conformation of the IgE Fc and create a discontinuous, multidomain epitope dominated by shape complementarity and networks of hydrogen bonds and salt bridges [19, 22] (Fig. 1B–C).

**Figure 1:**
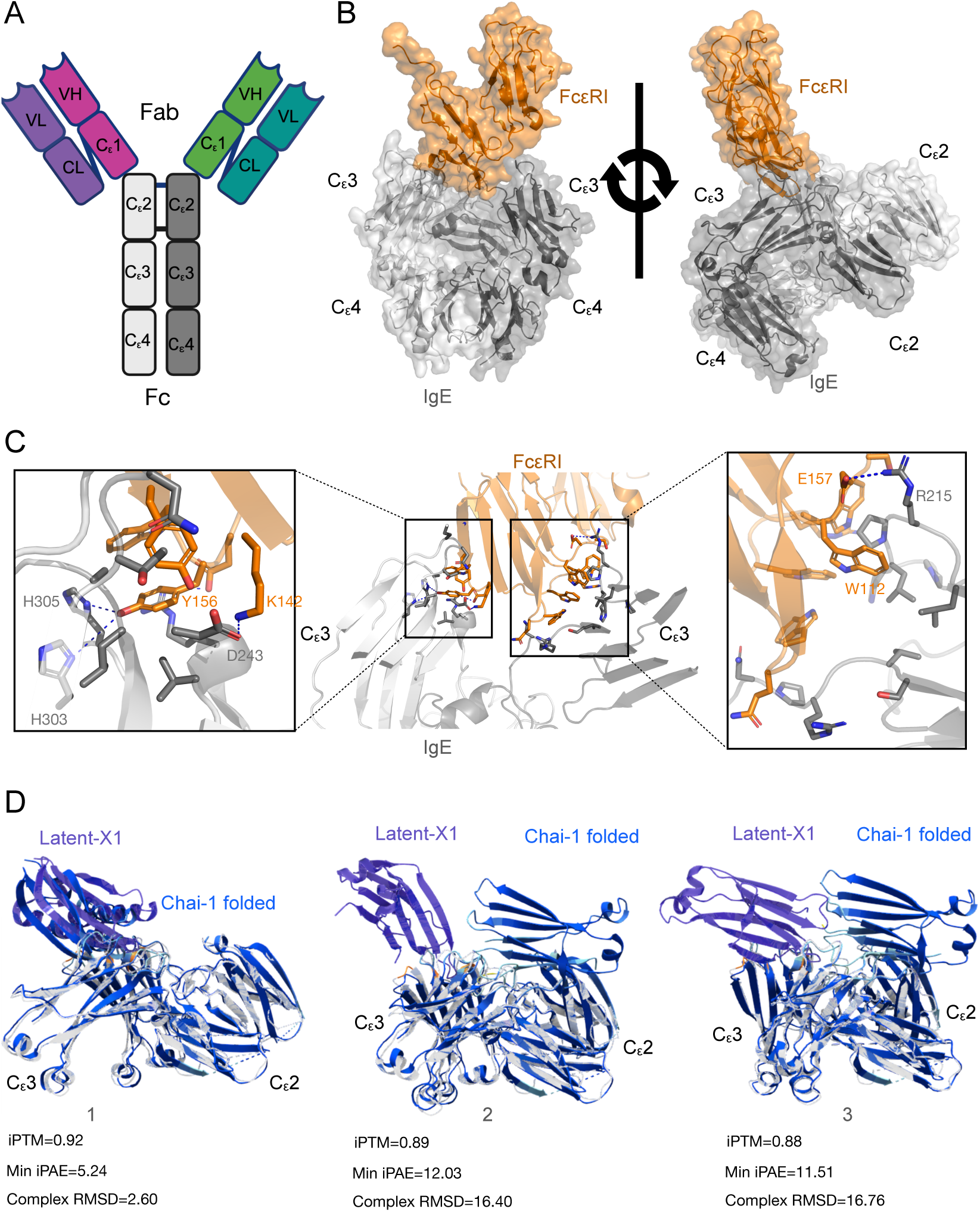
Defining IgE design prompts for targeting the Fc*ε*RI-binding site. (A) Domain organization of human IgE. (B) Cryo-EM structure of human IgE Fc (grey) bound to Fc*ε*RI (orange) (PDB: 8K7R), shown in two orientations. (C) Key interactions at the IgE–Fc*ε*RI interface highlighting hotspot residues on IgE involved in receptor engagement. (D) Representative top-ranked Latent-X1 designs (purple) are aligned with Chai-1 refolded complexes (blue), with accompanying model metrics (iPTM, min-iPAE, and complex RMSD in Å)

Importantly, both IgE Fc and Fc*ε*RI*α* are heavily glycosylated, and these glycans contribute to receptor binding, conformational stability, and signaling function [23–25]. The presence of dense and heterogeneous glycosylation further complicates binder design by introducing steric constraints and dynamic surface features that are not readily captured by standard protein-only modeling approaches. Together, this architecture underlies the sub-nanomolar affinity and exceptional stability of the IgE–Fc*ε*RI complex, enabling persistent mast-cell and basophil sensitization during allergic responses [16].

To initiate *de novo* binder design, we selected the cryo-EM structure of IgE Fc bound to Fc*ε*RI (PDB: 8K7R, 3.56 Å resolution), which captures the Fc region in a receptor-competent conformation [26]. A related structure (PDB: 8Z0T) was considered but excluded due to lower resolution and increased local disorder [27]. Full-length IgE Fc contains more than 300 residues per chain; when modeled as a homodimer, the total sequence exceeds 600 residues—surpassing the Latent-X1 512 context-length limit for joint binder–target design (Fig. 1D). To remain within these constraints while preserving the receptor-binding geometry, we isolated a truncated C*_ε_*2–C*_ε_*3 fragment that retains the full Fc*ε*RI-binding surface when reconstructed as a dimer.

We selected six interface residues spanning the asymmetric Fc*ε*RI-binding site as design hotspots and used binder lengths of 120 or 150 residues as initial prompts for Latent-X1–based design [13]. Across these runs, the model produced reasonable global-confidence scores (iPTM), indicating that the generated complexes displayed reasonable interfaces. However, two structure-accuracy metrics were uniformly poor: the min iPAE and the complex RMSD obtained after Chai-1 refolding. According to the Latent-X1 technical report, successful designs require approximately iPTM > 0.8, min iPAE < 1.0 Å, complex RMSD < 2.0 Å when assessed against the Chai-1 refolded complex [13]. In contrast, our best IgE-directed designs exceeded these thresholds by wide margins, with min iPAE values of 5–12 Å and complex RMSD values of 3–16 Å, despite acceptable iPTM (Fig. 1D). This pattern is characteristic of geometrically complex or discontinuous target epitopes: while the model can generate compact and internally plausible binders, it fails to produce complexes with low interfacial uncertainty or conformations that are consistent across orthogonal structure prediction models.

### Improving model generation through a single-domain seed epitope subsampling strategy

Despite extensive sampling, most Latent-X1 generations against the full Fc*ε*RI-binding site failed to produce high-quality complexes. Although the model positioned binders near the correct receptor-binding surface, the resulting designs showed poor agreement with Chai-1 refolding, yielding uniformly high min iPAE and large complex RMSD values (Fig. 2A). These results indicate that design against multichain, multidomain epitopes with inter-domain geometry remains challenging, leading to designs that appear locally plausible but fail to resolve accurate global interface geometry

**Figure 2:**
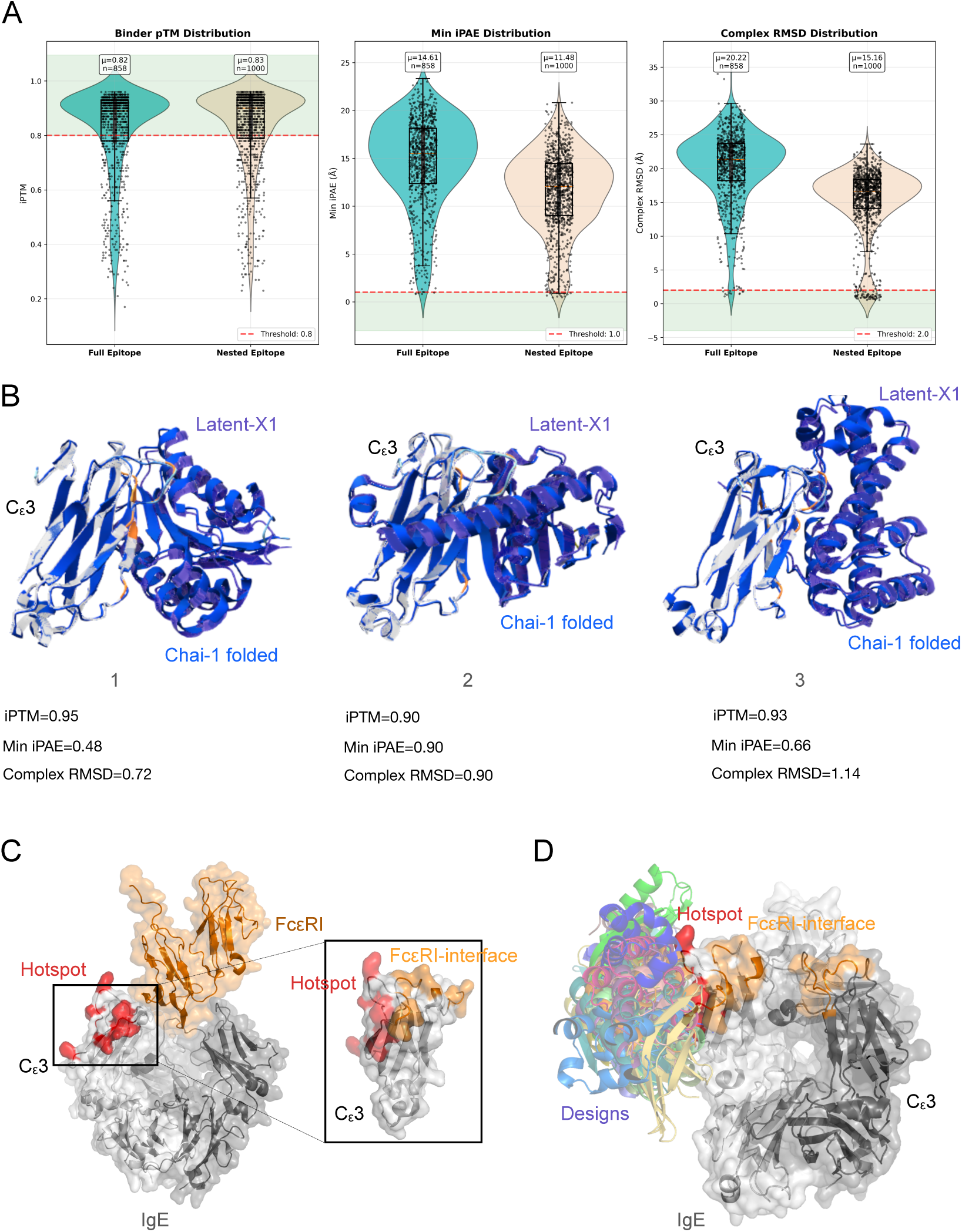
Nested epitope selection and binder design. (A) Violin plots comparing iPTM, min-iPAE, and complex RMSD for Latent X designs targeting the full Fc*ε*RI-binding interface (teal) or a restricted C*_ε_*3 seed epitope (tan). (B) Nested epitope designs show markedly improved interfacial accuracy and reduced geometric error after Chai-1 refolding based on model metrics (iPTM, min-iPAE, and complex RMSD in Å). (C) Structural schematic illustrating the nested epitope located adjacent to the Fc*ε*RI interface. Inset shows the C*ε*3 template with hotspot residues and the Fc*ε*RI binding site highlighted. (D) Superposition of top 10 Latent-X1 generations (purple) with the full-length IgE Fc structure, showing that several binders partially overlap with the Fc*ε*RI binding site while maintaining compatibility with the multidomain IgE architecture.

This motivated the development of a seed-epitope subsampling strategy focused initially on the C*ε*3 domain, which contains the majority of critical Fc*ε*RI-contact residues. We generated truncated C*ε*3 constructs and systematically scanned for local surface patches that (i) partially overlapped with the receptor interface or (ii) formed well-defined, solvent-accessible grooves with mixed polarity—features that typically support *de novo* binder accommodation. This approach immediately yielded substantial improvements in model quality relative to designs targeting the full discontinuous epitope (Fig. 2A). Whereas initial full-interface attempts produced min iPAE values in the 5–12 Å range and complex RMSD values of 6–13 Å after Chai-1 refolding, nested-epitope generations showed dramatically improved metrics. Across the top-performing design jobs, min iPAE values ranged from 1.36–1.77 Å and complex RMSD values from 1.01–1.26 Å, representing an order-of-magnitude improvement over previous attempts (Fig. 2A). This shift indicates that reducing geometric complexity enabled the model to generate binder–target poses that were consistent and better supported by Chai-1 refolding, thereby providing a viable foundation for wetlab success.

Guided by these results, we launched multiple Latent-X1 jobs targeting the best-performing nested epitope, exploring binder lengths of 100, 120, and 150 residues (Fig. 2B). Across these campaigns, we accumulated more than 600 top-ranked designs, each evaluated using three quantitative metrics. Visual inspection of high-scoring complexes revealed well-packed interfaces, appropriate stereochemistry, and no major atomic clashes, suggesting that many generated solutions were at least geometrically plausible.

To prioritize which sequences would advance to wet-lab screening, we developed a weighted scoring function that integrates all three metrics into a single ranking criterion:

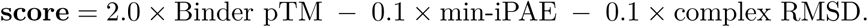

This weighting scheme reflects the different sensitivities of each metric: pTM acts as a global confidence indicator and thus receives the strongest weight, whereas min iPAE and complex RMSD provide structural-accuracy penalties that help deprioritize poses inconsistent with Chai-1 refolding. Because pTM varies over a narrow range (0.7–0.95) whereas iPAE and RMSD span much larger numerical scales, the chosen coefficients balance their relative contributions to yield a meaningful composite score.

Under this scheme, our nested-epitope generations produced composite scores ranging from 0.7–1.8—substantially better than scores obtained from full-epitope designs, which rarely exceeded 0.2.

This composite scoring approach allowed for borderline but potentially viable designs to be reconsidered rather than discarded by any single metric. By selecting representatives across this score spectrum, we ensured that both conservative and fringe candidates were sampled experimentally, ultimately enabling identification of productive binders that would have been missed by stricter, single-metric filtering.

Because our design prompt contained only a single IgE domain, the Latent-X1 model was not explicitly aware of the spatial context contributed by the remaining Fc domains or the second IgE chain. To ensure that the resulting complexes were compatible with the full multidomain IgE architecture, we performed detailed visual inspection of the top 40 ranked designs (Fig. 2C). This step verified that each binder could be superimposed onto the complete IgE Fc without steric clashes, domain interpenetration, or occlusion of structurally constrained regions, thereby confirming biological plausibility and compatibility with recombinant antigen formats used in downstream assays. All 40 designs passed this structural sanity check and were advanced to biolayer interferometry (BLI) validation.

### Rapid high-throughput Biolayer Interferometry (BLI) validation of miniproteins

To evaluate whether the *de novo* designed miniproteins bind human IgE Fc, we applied Onava’s rapid high-throughput BLI workflow, which enables sequence-to-sensorgram characterization in approximately seven days (Fig. 3A). This platform allows hundreds of designs to be evaluated in parallel, providing reproducible binding metrics suitable for model post-training, binder prioritization, and downstream functional studies such as epitope validation and competition assays.

**Figure 3:**
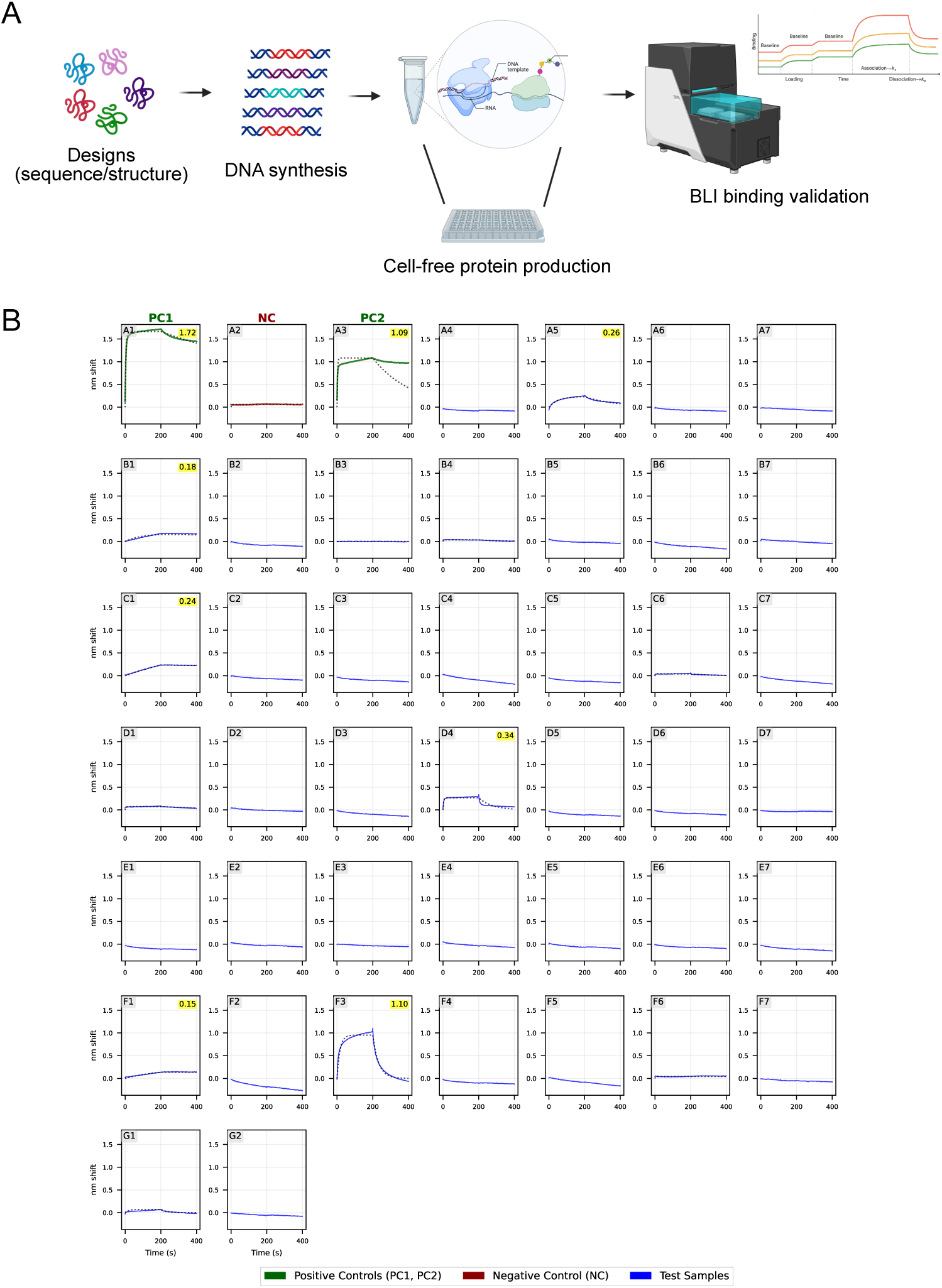
High-throughput BLI screening of IgE miniprotein designs. (A) Onava’s high-throughput binder validation workflow. (B) Sensorgrams for all tested miniproteins binding to human IgE Fc. Positive controls were: PC1, an omalizumab scFv in VH/VL orientation; and PC2, an omalizumab scFv in VL/VH orientation, both showing strong nm-shift responses. The negative control (NC) was trastuzumab scFv (VH/VL), which displayed minimal binding. Each panel shows the nm-shift (y-axis) over time (x-axis). Blue traces indicate designed IgE-binding candidates; green traces indicate positive controls; red traces indicate the negative control. Yellow labels indicate the maximal nm-shift observed for samples that exhibited detectable binding.

Each design was assessed for binding to recombinant human IgE Fc alongside two positive controls—PC1 and PC2, consisting of the omalizumab scFv in VH/VL and VL/VH orientations, respectively—and a negative control (NC), a trastuzumab scFv. Controls behaved as expected: both omalizumab formats produced strong nm-shift responses, while the trastuzumab control showed minimal binding (Fig. 3B). Designed miniproteins were tested under identical assay conditions, and their sensorgrams were plotted as nm shift over time, with yellow annotations indicating the maximum response for designs exhibiting signal above the NC baseline. Across the 40 designs screened, several candidates exhibited clear binding responses—though weaker than the omalizumab controls—confirming that the nested epitope strategy successfully produced the first generation of IgE-binding miniproteins (Fig. 3B).

We selected the top three binders—A5, D4, and F3—hereafter referred to as ONA-IgE-Rd1-1, ONA-IgE-Rd1-2, and ONA-IgE-Rd1-3, respectively—based on single-dose BLI responses exceeding 0.2 nm shift for progression to dose-dependent kinetic characterization (Fig. 4A). For these experiments, samples were immobilized onto BLI probes at matched loading levels and challenged with serial dilutions of human IgE Fc, starting at 1 *µ*M followed by three-fold serial dilutions. Omalizumab scFv (VH/VL orientation) was included as a positive control, while a trastuzumab scFv (VH/VL orientation) served as a negative control.

**Figure 4:**
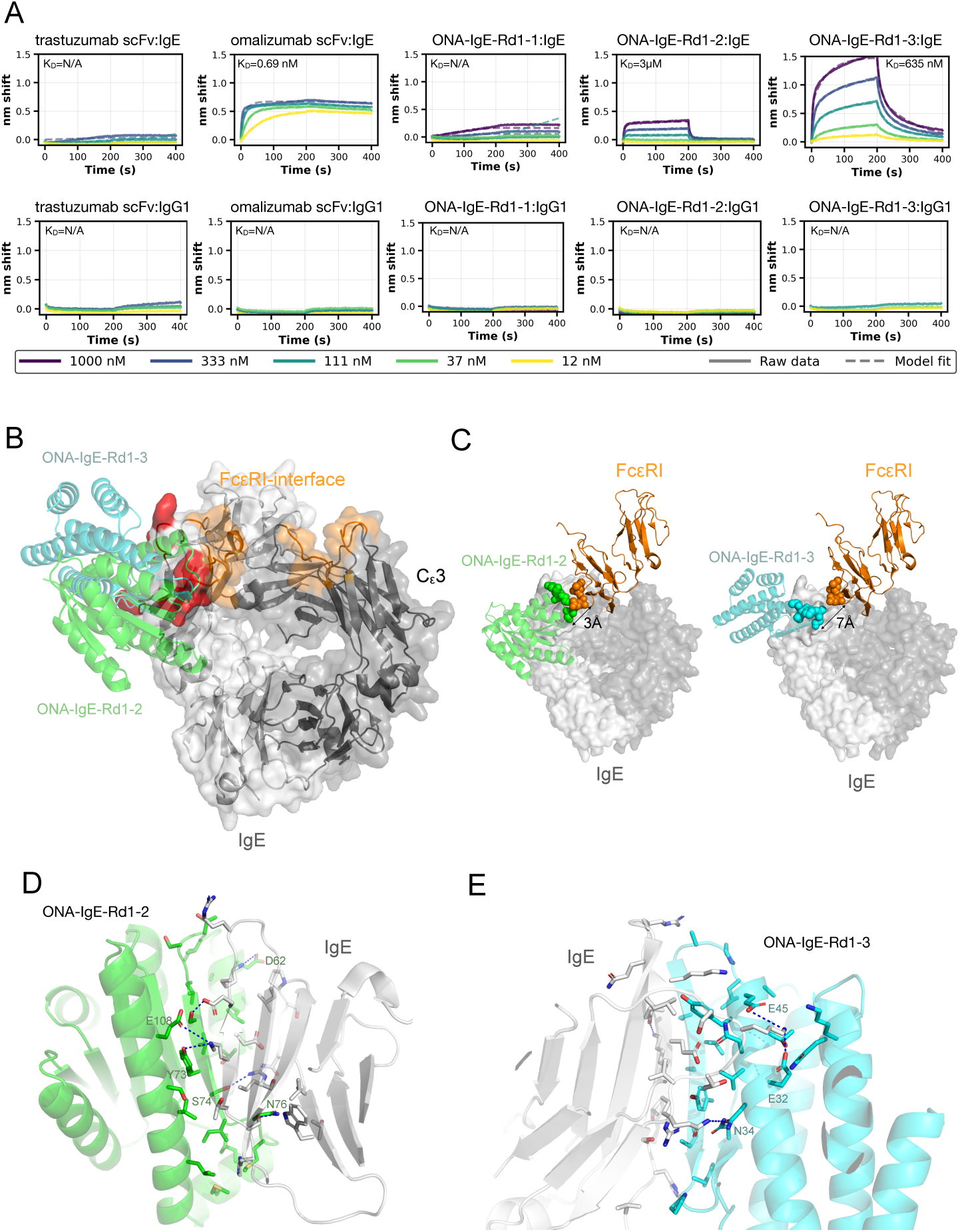
Biophysical validation and structural analysis of first-round IgE-binding miniproteins. (A) Dose-dependent BLI sensorgrams for validated miniprotein hits binding to the Fc region of human IgE (positive target) and human IgG1 Fc (negative target). Omalizumab scFv and trastuzumab scFv were included as positive and negative controls, respectively. IgE and IgG1 concentrations were tested as two-fold serial dilutions, as indicated below each sensorgram. (B) Alignment of Latent-X1 structural models for verified hits ONA-IgE-Rd1-2 (green) and ONA-IgE-Rd1-3 (cyan) onto their respective IgE binding interfaces. The Fc*ε*RI binding site is shown in orange, and the nested seed epitope used for design is highlighted in red. (C) Structural superposition of ONA-IgE-Rd1-2 and ONA-IgE-Rd1-3 with Fc*ε*RI, illustrating partial spatial overlap with the receptor-binding site. Distances between nearest-neighbor residues of the miniproteins and Fc*ε*RI are indicated for each complex. (D,E) Close-up views of the binding interfaces for ONA-IgE-Rd1-2 and ONA-IgE-Rd1-3, respectively, highlighting intermolecular *β*-sheet augmentation between the miniproteins and the IgE C*_ε_*3 domain. Key hydrogen bonding networks and stabilizing electrostatic interactions that contribute to IgE recognition are shown as dotted lines.

Dose-dependent kinetic analysis revealed that ONA-IgE-Rd1-2 and ONA-IgE-Rd1-3 bind human IgE Fc with dissociation constants (*K_D_*) of approximately 3 *µ*M and 635 nM, respectively. In contrast, ONA-IgE-Rd1-1 produced a weak response comparable to the trastuzumab negative control, indicating that this construct is likely a false positive. Similar weak or ambiguous signals were observed for a subset of designs in single-dose screening, underscoring the importance of secondary kinetic validation when working with proteins generated in cell-free expression systems, which can occasionally produce misleading signals due to partial misfolding or assay-related artifacts. To further assess specificity, human IgG1 Fc was included as an orthogonal control antigen; no binding was observed for any of the designs, confirming that the measured interactions are specific to IgE Fc.

We next analyzed the binding interfaces of the validated hits ONA-IgE-Rd1-2 and ONA-IgE-Rd1-3 to identify structural features and residues contributing to IgE recognition. Superposition of both miniproteins onto the full-length IgE Fc structure showed that they bind at positions nested adjacent to the Fc*ε*RI interface, consistent with the epitope defined during the design process (Fig. 4B). Despite targeting the same epitope region, the two binders adopt distinct architectures: both are mixed *α/β* proteins but share limited sequence similarity and only modest structural resemblance, highlighting the scaffold diversity accessible through Latent-X1 generative design.

Spatial analysis of the modeled complexes revealed minimum distances of approximately 3 Å for ONA-IgE-Rd1-2 and 7 Å for ONA-IgE-Rd1-3 relative to Fc*ε*RI, suggesting that both binders occupy positions sufficiently proximal to interfere with receptor engagement (Fig. 4C). From a structural perspective, these binding modes imply that miniproteins could simultaneously engage independent sites on each C*_ε_*3 domain within the IgE Fc homodimer, potentially enhancing competition with Fc*ε*RI.

Detailed inspection of the interfaces showed that, in both complexes, binding is dominated by *β*-sheet augmentation: *β*-strands from the designed miniproteins extend and pair with native *β*-strands of the IgE C*_ε_*3 domain, forming a continuous intermolecular sheet. Additional stabilizing interactions arise from a neighboring IgE *α*-helix, which contributes auxiliary contacts that reinforce the interface (Fig. 4D–E). Across both designs, we observe networks of hydrogen bonds and salt bridges that collectively stabilize the complexes and rationalize the observed binding affinities.

### Structure-Guided Epitope Expansion Improves Competition with Fc***ε***RI

Our overarching design objective was to generate compact miniprotein binders capable of directly competing with Fc*ε*RI for binding to IgE. While round-one designs yielded validated IgE-binding miniproteins, structural analysis revealed that these hits engaged only a subset of the Fc*ε*RI-binding interface and were therefore unlikely to fully occlude receptor engagement. To address this limitation, we pursued a second round of structure-guided design focused on expanding the binder interaction surface into regions of the IgE C*_ε_*3 domain directly involved in Fc*ε*RI recognition.

Building on the successful seed-epitope strategy established in round one, we systematically explored controlled expansion of the target epitope within the IgE C*_ε_*3 domain toward the Fc*ε*RI-binding site. Accessible surface area and interface analysis of the IgE–Fc*ε*RI complex identified several solvent-accessible residues, structurally stable positions that contribute to the receptor-binding interface and could plausibly be incorporated into an expanded binder epitope without perturbing the local fold (Fig. 5A). These residues lie immediately adjacent to the round-one seed epitope and collectively define a contiguous interaction surface suitable for second-round design (Fig. 5A).

**Figure 5:**
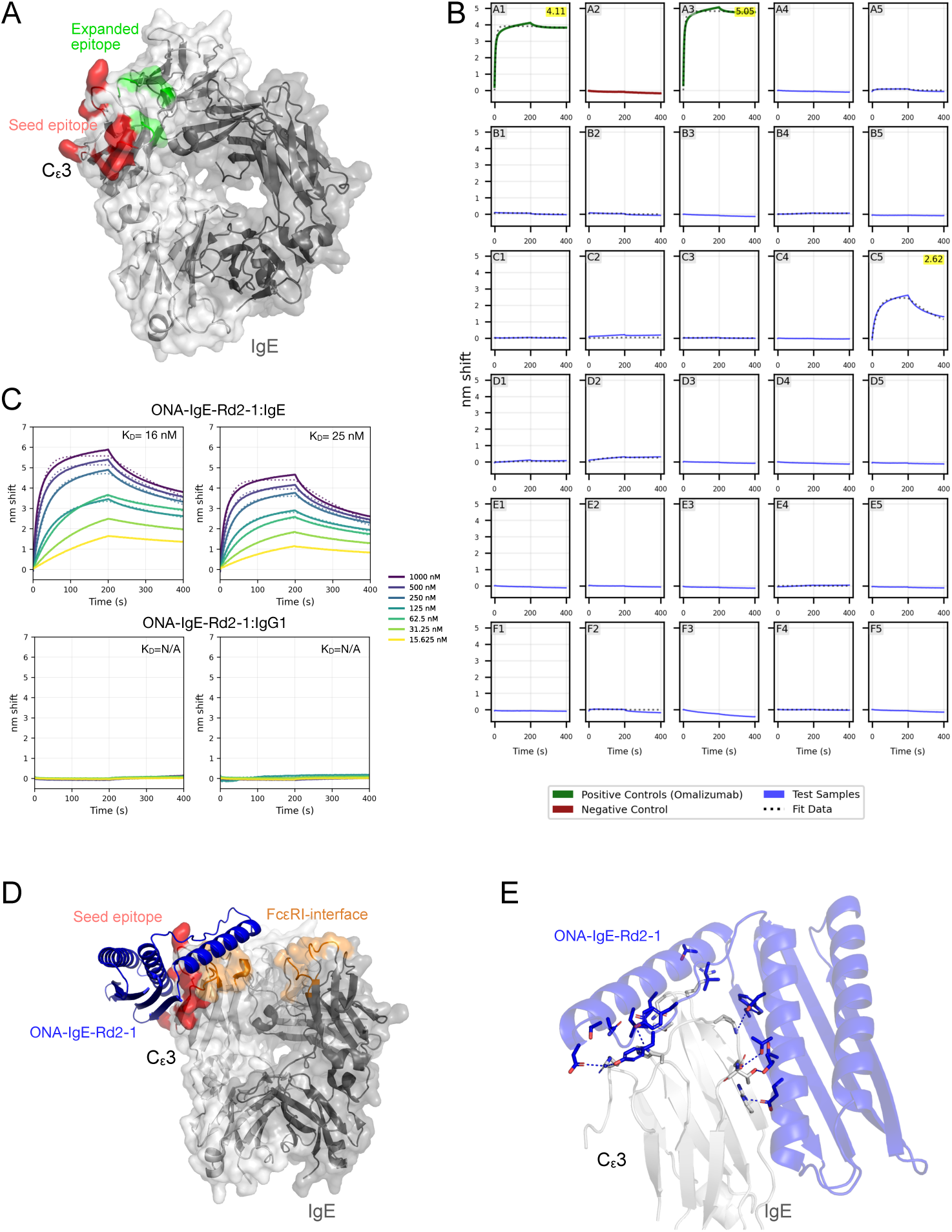
Characterization of a round-two IgE-binding miniprotein obtained via epitope expansion. (A) Structural representation of the expanded epitope on IgE. The Fc*ε*RI-binding interface that comprise the expanded epitope is shown in green, and the nested seed epitope used to guide second-round design is highlighted in red. (B) Rapid BLI screening of the top 27 miniprotein designs generated in round two. Controls are as described in Figure 3B. (C) Dose-dependent kinetic validation of ONA-IgE-Rd2-1, showing replicate BLI sensorgrams for binding to human IgE Fc and the negative control antigen IgG1 Fc. (D) Structural superposition of ONA-IgE-Rd2-1 onto the full-length IgE Fc structure, illustrating extensive overlap with the Fc*ε*RI-binding interface. (E) Close-up views of the binding interface between ONA-IgE-Rd2-1 and the IgE C*_ε_*3 domain, highlighting intermolecular *β*-sheet augmentation and key residues contributing to IgE recognition. Hydrogen-bonding networks and stabilizing electrostatic interactions are indicated by dotted lines.

Using this expanded epitope definition, we initiated multiple Latent-X1 design batches, generating approximately 400 candidate binders. Designs were evaluated and ranked using the composite scoring framework described above, integrating binder pTM, min-iPAE, and complex RMSD, followed by visual inspection of interface geometry and packing. From this pool, the top 27 designs were selected for experimental screening and evaluated using our rapid seven-day BLI workflow alongside the same positive and negative controls employed in round-one screening. As expected, omalizumab scFv controls produced robust binding responses, while the majority of second-round designs exhibited baseline signals comparable to the trastuzumab scFv negative control (Fig. 5B).

Notably, a single design located at position F3 in the screening layout displayed a strong binding response, with a single-dose apparent *K_D_* of approximately 70 nM. This candidate was advanced for secondary, dose-dependent kinetic characterization and is hereafter referred to as ONA-IgE-Rd2-1. Across replicate experiments, ONA-IgE-Rd2-1 bound human IgE Fc with an average equilibrium dissociation constant of 19 nM, representing an approximately 30-fold improvement in affinity relative to the strongest round-one hit (Fig. 5C). Importantly, ONA-IgE-Rd2-1 showed no detectable binding to the negative control antigen human IgG1 Fc, confirming specificity for IgE and excluding nonspecific Fc interactions.

To understand the structural basis for the observed affinity improvement and enhanced competitive behavior, we examined the predicted complex structure of ONA-IgE-Rd2-1 in the context of full-length IgE.Superposition of the design model onto the IgE Fc structure revealed that ONA-IgE-Rd2-1 binds precisely at the intended expanded epitope, forming extensive overlap with the Fc*ε*RI-binding interface within a single C*_ε_*3 domain (Fig. 5D). This degree of interface overlap strongly suggests that ONA-IgE-Rd2-1 directly competes with Fc*ε*RI for binding, thereby fulfilling the primary objective of the epitope-expansion strategy.

At the molecular level, ONA-IgE-Rd2-1 retains the same fundamental binding topology observed in round-one hits, with the interaction dominated by *β*-sheet augmentation, in which *β*-strands from the miniprotein extend and pair with native *β*-strands of the IgE C*_ε_*3 domain (Fig. 5E). However, compared to round-one designs, the expanded interface of ONA-IgE-Rd2-1 displays increased hydrophobic surface burial and improved shape complementarity, accompanied by a modest reduction in the number of polar and electrostatic contacts. This redistribution of interaction types likely stabilizes the complex by enhancing packing efficiency and reducing solvent exposure, providing a structural explanation for the observed gain in affinity.

Together, these results demonstrate that structure-guided epitope expansion can be used to iteratively refine *de novo* miniprotein binders toward functional competition with a native receptor. By progressively extending a tractable seed epitope into a larger, biologically relevant binding interface, we were able to convert weak binders into higher affinity competitors of Fc*ε*RI, underscoring the power of this strategy for targeting complex, multidomain protein interfaces.

### Alanine-scanning mutagenesis validates the designed binding interfaces

To further assess the specificity and structural accuracy of both round-1 and round-2 hits, we performed systematic alanine-scanning mutagenesis of the miniprotein binding interfaces. For each Latent-X1 design model, residues within 5 Å of the IgE C*_ε_*3 domain were identified as interface positions and individually substituted with alanine (Fig. 6). All variants were codon-optimized using our internal protein-engineering pipeline and evaluated using the same rapid BLI workflow employed for primary binder validation.

**Figure 6:**
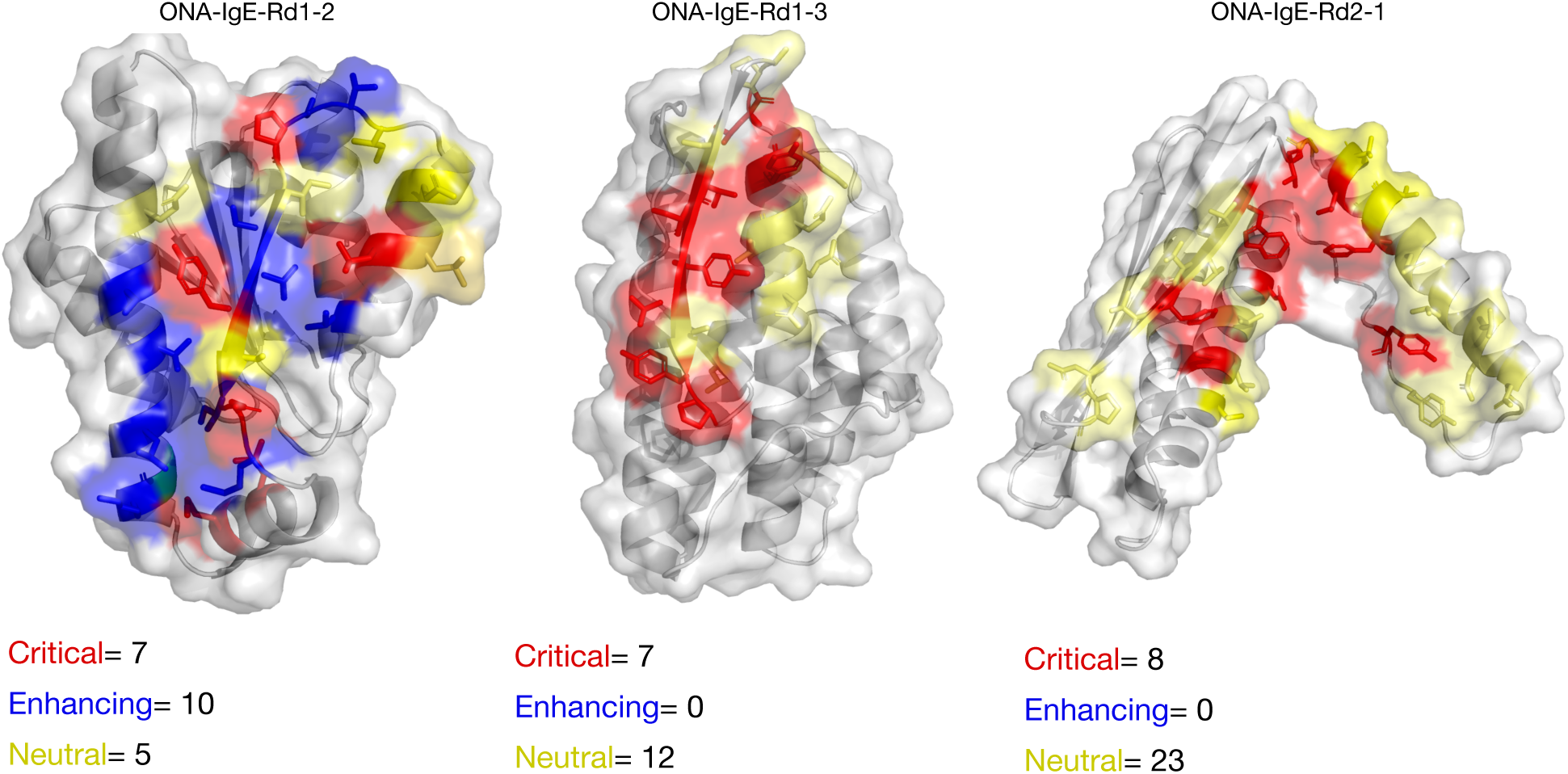
Alanine-scanning mutagenesis maps functional binding interfaces of miniprotein hits. Residues at the binding interfaces of ONA-IgE-Rd1-2, ONA-IgE-Rd1-3, and ONA-IgE-Rd2-1 are colored based on the impact of alanine substitution on IgE binding.

Mutant variants were classified based on their effect on IgE binding: substitutions resulting in a > 10*×* loss of binding affinity were scored as deleterious, substitutions with effects within a 10*×* window were considered neutral, and substitutions yielding a > 10*×* improvement in affinity were classified as affinity-enhancing. Across ONA-IgE-Rd1-2, ONA-IgE-Rd1-3, and ONA-IgE-Rd2-1, we observed that more than seven interface residues per binder produced substantial reductions in IgE binding upon alanine substitution, indicating that binding is distributed across a well-defined and cooperative interface rather than dominated by a single hotspot (Fig. 6).

For the round-1 binders ONA-IgE-Rd1-2 and ONA-IgE-Rd1-3, the strongest deleterious effects localized to residues positioned on *β*-strands involved in *β*-sheet augmentation with the IgE C*_ε_*3 domain, consistent with the structural models. Notably, in ONA-IgE-Rd1-2, several alanine substitutions resulted in greater than 10*×* affinity improvements, bringing binding into the 100–200 nM range and highlighting the potential for rapid affinity maturation through a small number of focused mutations. In contrast, non-deleterious mutations in ONA-IgE-Rd1-3 and the round-2 binder ONA-IgE-Rd2-1 yielded more modest improvements, with a maximum observed enhancement of approximately five-fold, suggesting that further gains in affinity may require combinatorial mutations and exploitation of epistatic effects (Fig. 6).

For ONA-IgE-Rd2-1, the most critical residues clustered within the deeply buried portion of the interface (Fig. 6). In particular, alanine substitution of a residue forming a buried salt bridge with IgE residue K440 resulted in complete loss of binding, underscoring the importance of electrostatic stabilization at this site. Collectively, these mutational data delineate the key residues and structural motifs required for IgE recognition and provide strong experimental validation that the designed interfaces mediate binding in all validated hits.

### Polyspecificity profiling of IgE-binding miniproteins

Although no binding was detected to IgG1 Fc in our BLI assays, we next sought to rigorously assess the specificity and polyspecificity liabilities of IgE-binding miniprotein hits. This analysis was motivated by the fact that these binders engage IgE Fc through non-canonical, *de novo*–designed interfaces that differ substantially from those used by natural antibodies. To this end, we performed BVP-ELISA on all validated hits and included two high-affinity miniproteins from prior published studies—CbAgo_b2 and GDM_VEGFA_79—as reference controls [6, 10]. These comparators were selected to benchmark the polyspecificity behavior of our designs against well-known *de novo* miniproteins reported in high-impact literature and often assumed to possess favorable developability properties. Antibody-derived controls routinely used in polyreactivity assays were included in parallel to provide context relative to established monoclonal benchmarks.

To decouple intrinsic molecular polyspecificity from artifacts introduced by protein production, we evaluated both His-tagged and Fc-tagged formats for all miniproteins. His-tagged constructs were expressed using our cell-free expression system and purified via Ni–NTA affinity chromatography (Supplementary Data), while Fc-tagged variants were expressed in Expi-CHO cells and purified using Protein A affinity chromatography followed by size-exclusion chromatography (SEC). All proteins were obtained at high overall purity; however, SEC analysis revealed that several samples exhibited reduced main-peak fractions, indicative of partial heterogeneity. This parallel production strategy enabled direct comparison across expression platforms, purification methods, and molecular formats, allowing us to isolate the contribution of avidity to observed binding behavior.

For BVP-ELISA, all samples were normalized and tested at three concentrations (350 nM, 70 nM, and 14 nM). His-tagged antibody scFv controls produced in cell-free reactions were included alongside monoclonal antibody controls expressed in Expi-CHO cells. BVP-ELISA scores were calculated from triplicate measurements and normalized to their respective controls. These experiments revealed several notable trends (Fig. 7A). In the His-tagged format, ONA-IgE-Rd1-2 and ONA-IgE-Rd2-1 displayed low polyspecificity scores comparable to highly specific antibody controls such as trastuzumab and daclizumab scFvs. The VEGF-binding miniprotein control GDM_VEGFA_79 similarly exhibited low off-target reactivity. In contrast, ONA-IgE-Rd1-3 and the published miniprotein CbAgo_b2 showed elevated polyspecificity scores at the highest concentration (350 nM), though still below those observed for the highly polyreactive antibody control bococizumab (Fig. 7A).

**Figure 7:**
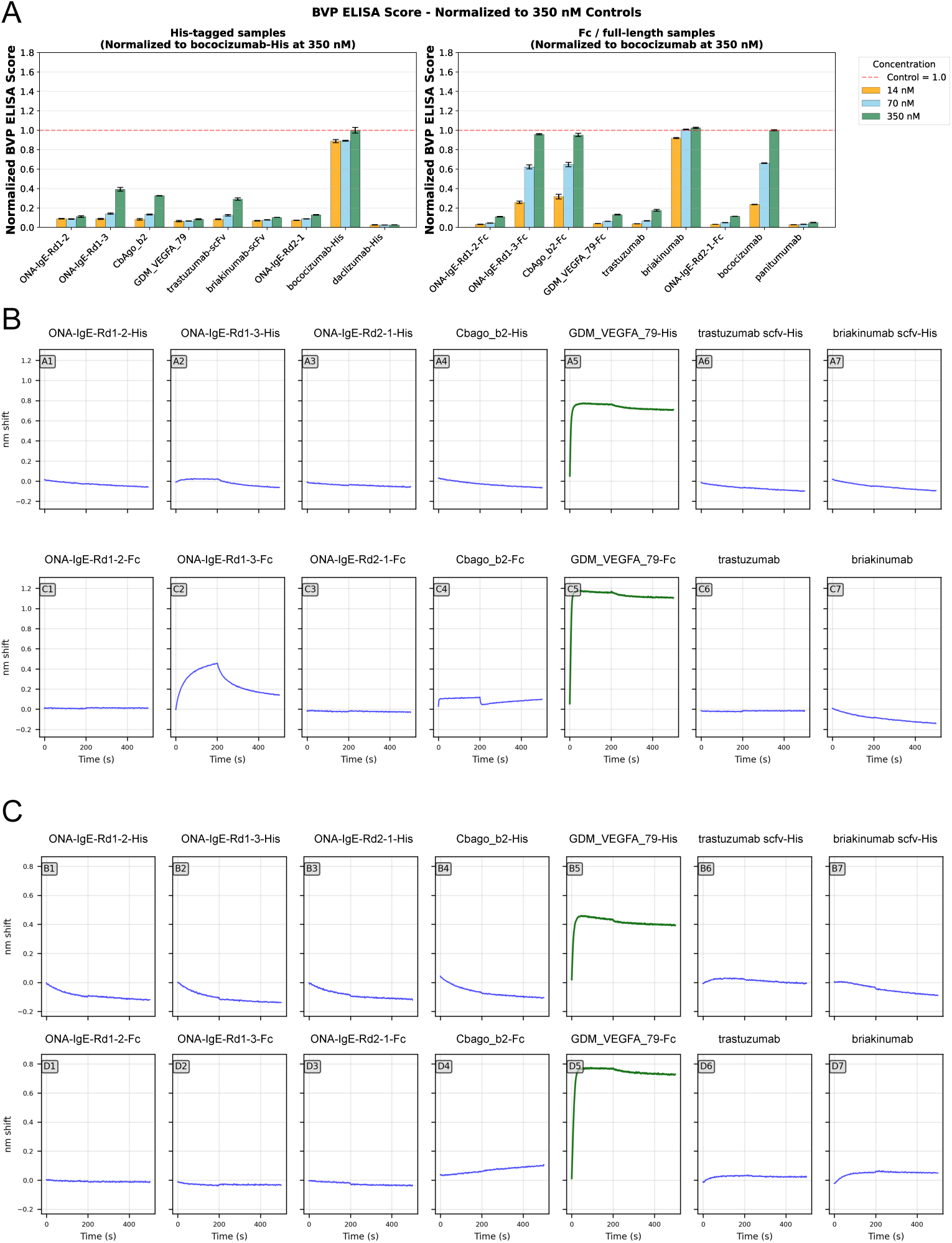
Polyspecificity and off-target binding analysis of IgE-binding miniproteins. (A) BVP-ELISA polyspecificity scores for His-tagged (left) and Fc-fusion (right) formats tested at 14 nM, 70 nM, and 350 nM. Scores are shown as mean *±* SD (n=3) and normalized to the highly polyreactive monoclonal antibody control bococizumab. Antibody-derived specificity controls (trastuzumab, daclizumab, and panitumumab) and published miniprotein benchmarks (CbAgo_b2 and GDM_VEGFA_79) are included for comparison. (B,C) BLI sensorgrams evaluating off-target binding to VEGF165 (B) and VEGF110 (C). The VEGF-binding miniprotein GDM_VEGFA_79 (green) serves as a positive control and displays robust binding to both VEGF isoforms.

Strikingly, the Fc-tagged format substantially altered polyspecificity behavior. While ONA-IgE-Rd1-2-Fc, ONA-IgE-Rd2-1-Fc, and GDM_VEGFA_79-Fc maintained low off-target BVP signals comparable to trastuzumab and panitumumab monoclonal antibodies, both ONA-IgE-Rd1-3-Fc and CbAgo_b2-Fc exhibited a pronounced increase in polyspecificity (Fig. 7A). These Fc fusions reached BVP-ELISA scores similar to those of bococizumab and briakinumab, two monoclonal antibodies well known for their polyreactive liabilities. Notably, this potentiation of off-target binding was evident at concentrations as low as 70 nM.

We further evaluated polyspecific binding of His-tagged and Fc-tagged miniprotein samples using BLI against the multidomain antigen VEGF165 (Fig. 7B). VEGF165 was selected because its shorter isoform, VEGF110, lacks the C-terminal heparin-binding domain, enabling discrimination between domain-specific off-target interactions and nonspecific binding. In this experiment, the VEGF-binding miniprotein GDM_VEGFA_79—which exhibited low polyreactivity in BVP-ELISA—served as a positive control binder, while trastuzumab was included as a negative control. Both controls behaved as expected, validating the assay conditions. Consistent with the BVP-ELISA results, most His-tagged miniprotein samples showed no detectable binding to VEGF165 by BLI. As expected, the positive control GDM_VEGFA_79 exhibited strong binding, while trastuzumab showed no response. A weak but reproducible VEGF165 binding signal was observed for the polyreactive variant ONA-IgE-Rd1-3-His, in agreement with its elevated polyspecificity score at higher concentrations in BVP-ELISA (Fig. 7B).

In contrast, Fc-tagged formats revealed a pronounced avidity-dependent effect. ONA-IgE-Rd1-3-Fc displayed significant binding to VEGF165, and a weaker but detectable interaction was observed for CbAgo_b2-Fc (rmax 0.18) (Fig. 7B). These trends closely mirrored the BVP-ELISA results. Importantly, no binding was detected for any sample against VEGF110, except the VEGF-binding miniprotein control GDM_VEGFA_79, indicating that the observed off-target interactions for ONA-IgE-Rd1-3-Fc and CbAgo_b2-Fc arise from interactions with the C-terminal domain unique to VEGF165 rather than from nonspecific surface adsorption(Fig. 7C).

Collectively, these results demonstrate that polyspecificity in *de novo* miniproteins can be strongly amplified by avidity and molecular format, particularly upon Fc fusion. Importantly, this behavior is not unique to our designs but is also observed in previously published high-affinity miniproteins, underscoring a broader and underappreciated challenge in the field. These findings highlight a critical gap between *in silico* design success and downstream developability, and emphasize the need for generative protein design strategies to explicitly account for polyspecificity and avidity-driven off-target interactions early in the design process rather than relying solely on affinity-based screening.

### IgE-binding miniproteins compete with the native Fc***ε***RI receptor

To evaluate the functional consequence of our designs, we assessed whether IgE-binding miniproteins could compete with the native Fc*ε*RI receptor for binding to IgE. Fc*ε*RI engages IgE with sub-nanomolar affinity, and thus any effective competition by miniproteins would be expected to require either substantial affinity maturation or favorable steric and geometric exclusion. We therefore explored multiple BLI-based competition formats to probe competitive binding under different molecular orientations and loading strategies.

We first established baseline binding affinities for Fc-tagged miniproteins using dose-dependent kinetic titrations in which IgE Fc was immobilized on biosensors via His-tag capture and Fc-tagged miniproteins were introduced as analytes at concentrations starting at 2 *µ*M with two-fold serial dilution (Fig. 8A). Under these conditions, ONA-IgE-Rd1-2, ONA-IgE-Rd1-3, and ONA-IgE-Rd2-1 bound IgE with apparent dissociation constants of 1.2 *µ*M, 285 nM, and 60 nM, respectively. Relative to their His-tagged counterparts, Fc-tagged ONA-IgE-Rd1-2 and ONA-IgE-Rd1-3 showed approximately three-fold improvements in affinity, consistent with avidity effects, whereas ONA-IgE-Rd2-1 exhibited a modest decrease in apparent affinity driven by a slower association rate.

**Figure 8:**
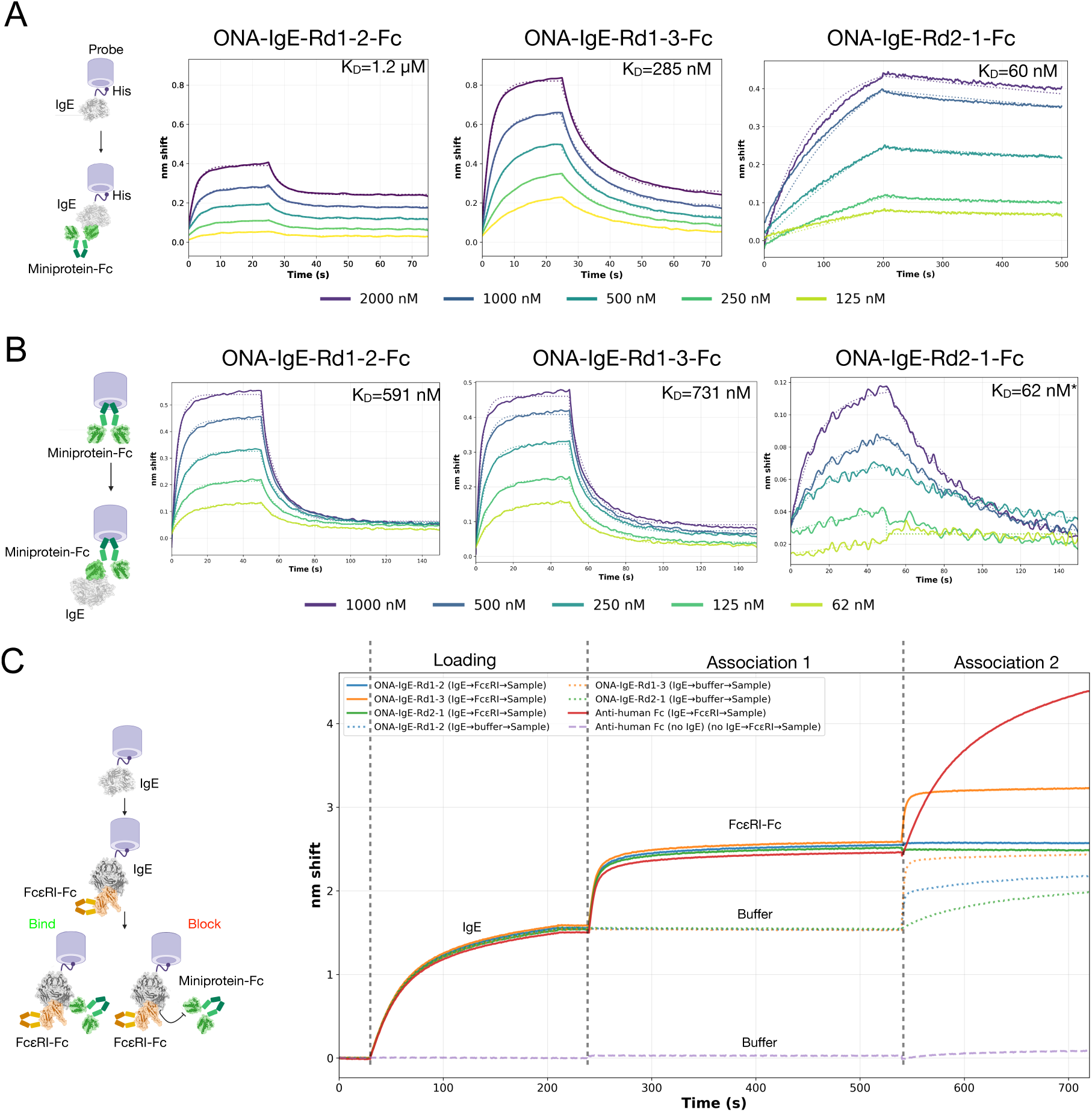
Competition BLI validates overlap between IgE-binding miniproteins and the Fc*ε*RI binding interface. A) Dose-dependent kinetic titrations of Fc-tagged miniproteins in a configuration where IgE Fc is immobilized on the biosensor and Fc-tagged miniproteins are measured as analytes. B) Reciprocal titrations with Fc-tagged miniproteins immobilized on the biosensor and IgE Fc measured as the analyte. C) Three-step tandem competition assay in which IgE-His is first immobilized, Fc*ε*RI-Fc (or buffer) is bound in Association 1, and Fc-tagged miniproteins are tested in Association 2. Solid traces indicate the competition format (Fc*ε*RI pre-bound), while dotted traces omit Fc*ε*RI to report direct IgE binding. The no-IgE condition reports background. Under Fc*ε*RI pre-binding, ONA-IgE-Rd1-2 and ONA-IgE-Rd2-1 are blocked, whereas ONA-IgE-Rd1-3 retains binding, consistent with a non-overlapping or alternative binding mode. Anti-human IgG1 Fc used as positive control of non-competitve binding to the IgE-Fc*ε*RI complex

We next examined binding in the reciprocal orientation, immobilizing Fc-tagged miniproteins on Protein A biosensors and measuring IgE Fc as the analyte (Fig. 8B). In this configuration, ONA-IgE-Rd1-2 and ONA-IgE-Rd1-3 displayed clear binding signals with apparent dissociation constants of 591 nM and 731 nM, respectively. In contrast, ONA-IgE-Rd2-1 produced a weaker and noisier response, yielding a fitted *K_D_* of approximately 62 nM but with signal amplitudes close to background. We speculate that this orientation introduces steric constraints for ONA-IgE-Rd2-1, potentially due to linker geometry or restricted accessibility of the binding interface, suggesting that linker optimization may be required for robust binding in Fc-immobilized formats.

Based on the superior signal quality observed with IgE immobilized via His-tag capture, we proceeded to assess direct competition between Fc*ε*RI and miniproteins using a three-step tandem BLI competition assay (Fig. 8C). In this format, IgE Fc was first immobilized to saturation, followed by association with Fc*ε*RI–IgG1 Fc to occupy the receptor-binding interface or buffer as a control. In the final association step, Fc-tagged miniproteins or an anti-human IgG1 Fc antibody were introduced to evaluate binding to IgE under receptor-saturated or receptor-free conditions. The anti-human IgG1 antibody served as a positive control, yielding strong binding signals consistent with recognition IgE–Fc*ε*RI complex via IgG1 Fc tag on Fc*ε*RI.

When Fc*ε*RI was omitted in the first association step, all miniproteins exhibited robust binding in the final association phase, confirming their ability to bind immobilized IgE Fc. In contrast, pre-saturation of IgE with Fc*ε*RI resulted in complete blockade of binding for ONA-IgE-Rd1-2 and ONA-IgE-Rd2-1—two designs that also displayed low polyspecificity in BVP-ELISA assays (Fig. 8C). Notably, ONA-IgE-Rd1-3 retained strong binding even in the presence of Fc*ε*RI, with signal amplitudes comparable to those observed in the buffer control condition. This behavior is consistent with either non-competitive binding, as structural models place the closest distance to Fc*ε*RI at 7 Å, or engagement of alternative IgE epitopes enabled by increased polyspecificity. Importantly, ONA-IgE-Rd1-3 does not bind Fc*ε*RI directly in control experiments, indicating that observed interactions are mediated exclusively through IgE Fc.

Collectively, these competition studies demonstrate that two of the three validated miniprotein designs achieve the intended design objective of binding to the Fc*ε*RI-binding interface on IgE. However, despite clear competitive exclusion in vitro, the measured affinities remain insufficient to outcompete Fc*ε*RI under physiological conditions, highlighting the need for further affinity maturation and interface optimization.

## Discussion

Designing *de novo* binders against multidomain protein interfaces remains a central challenge in generative protein engineering. Although recent generative models have demonstrated impressive performance on compact, single-domain targets, their effectiveness degrades when applied to composite epitopes that span multiple domains or depend on precise inter-domain geometry. In this study, we show that these limitations are not intrinsic barriers, but can be overcome through an iterative, structure-guided epitope expansion strategy tightly coupled to rapid experimental validation.

A key finding is that design against the full Fc*ε*RI-binding interface of IgE consistently failed, despite extensive sampling. These failures were not apparent from *in silico* scores alone and only became evident upon refolding and experimental testing. By contrast, restricting the initial design space to a geometrically tractable seed epitope on the C*_ε_*3 domain yielded consistent models and experimentally validated binders. These first-round hits were not endpoints, but anchors for controlled epitope expansion into regions directly involved in Fc*ε*RI recognition. This stepwise approach enabled progressive increases in interface overlap and affinity, ultimately producing a double-digit nanomolar binder that engages the intended receptor-binding surface.

An additional and unexpected outcome of this work is that the *de novo* miniproteins engage an IgE epitope that is distinct from those observed in previously solved IgE–antibody or IgE–receptor complexes (Fig. 9A). Structural alignment of all available IgE-bound structures reveals two dominant clusters of antibody binding on the IgE Fc, whereas the designed miniproteins occupy a spatially separate surface with minimal overlap with these canonical epitopes. Furthermore, this binding mode does not overlap with the CD23 interaction surface, suggesting intrinsic receptor selectivity that is advantageous for therapeutic modulation of IgE (Fig. 9B). This binding mode emerges naturally from the nested epitope expansion strategy, which biases generation toward geometrically accessible but underexplored regions of the IgE surface rather than recapitulating immunologically selected binding sites.

**Figure 9:**
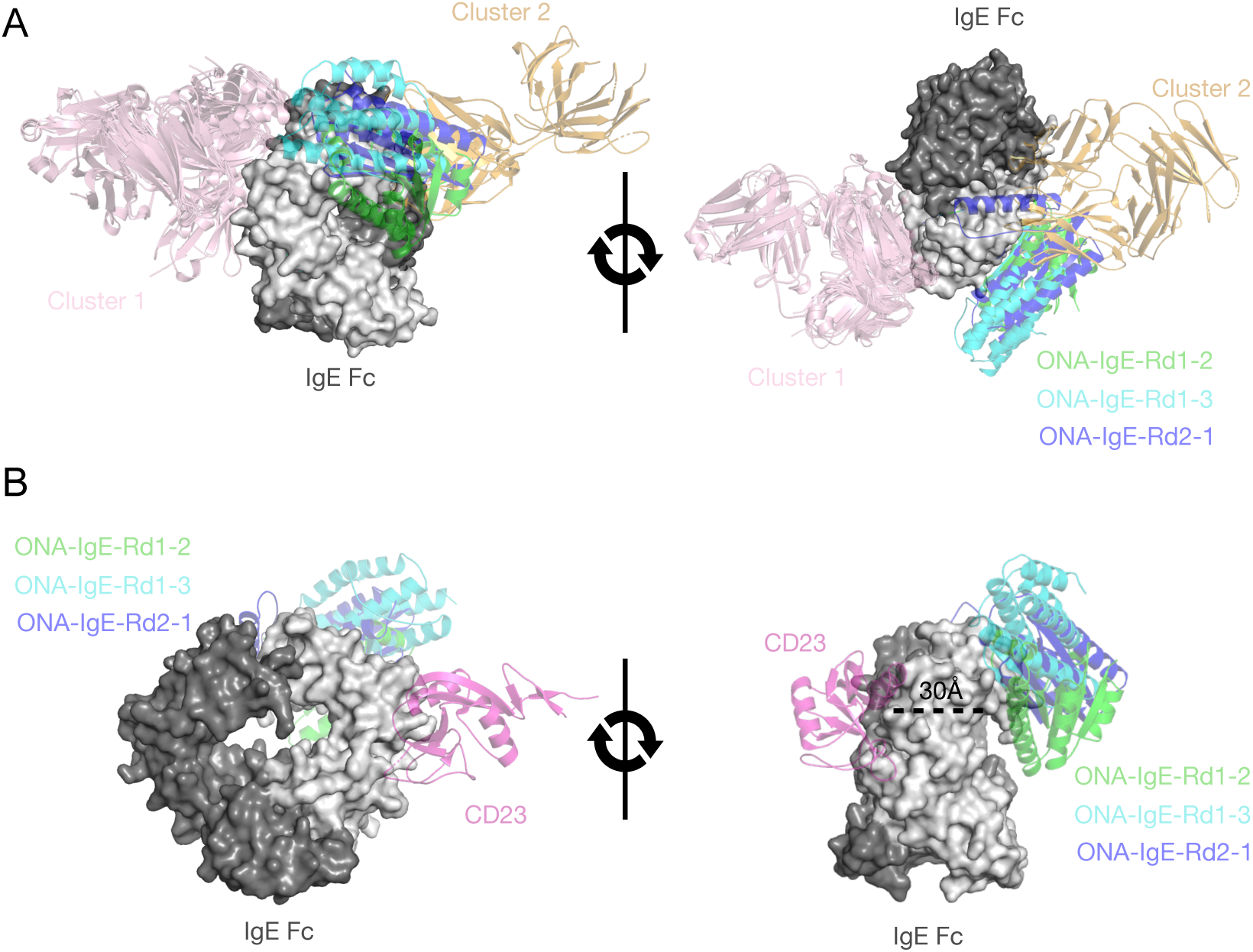
Designed IgE-binding miniproteins engage a distinct epitope on IgE Fc. (A) Structural alignment of all available IgE-bound complexes deposited in the PDB reveals two dominant clusters of antibody binding epitopes on the IgE Fc. Miniproteins cluster to a unique binding site that slightly overlaps with cluster 2. Cluster 1 maps to the omalizumab epitope, while cluster 2 aligns with the ligelizumab epitope (B) Structural superposition shows that the miniprotein binding interfaces are spatially distinct from the CD23-binding site, with the closest interfacial distance greater than 30 Å.

Engagement of a non-canonical epitope has important functional implications for modulating IgE–Fc*ε*RI interactions. The IgE–Fc*ε*RI complex is characterized by exceptionally high affinity and long receptor occupancy on mast cells and basophils, rendering direct competitive displacement intrinsically difficult once complexes are formed [28,29]. Consistent with this constraint, the affinities of the present designs are insufficient to displace Fc*ε*RI from pre-assembled IgE–receptor complexes. However, two binders exhibit partial overlap with the Fc*ε*RI-binding interface and compete with receptor binding, consistent with the intended design geometry. With additional affinity maturation, these miniproteins provide a foundation for probing the therapeutic potential of a non-canonical IgE epitope that has remained inaccessible to conventional antibody-based approaches.

These observations suggest a plausible path toward *facilitated dissociation* mechanisms, in which binding to an adjacent or partially overlapping epitope destabilizes the IgE–Fc*ε*RI complex rather than competing directly at equilibrium. Such mechanisms have been demonstrated using disruptive antibodies and engineered protein inhibitors that actively accelerate IgE–receptor dissociation and desensitize allergic effector cells [28, 30, 31]. Facilitated dissociation is particularly attractive in IgE biology because therapeutic intervention typically occurs after Fc*ε*RI is already occupied, a regime in which conventional competitive antibodies are sterically blocked and kinetically disadvantaged. By engaging a distinct but structurally coupled surface, targeted *de novo* binders may enable indirect destabilization of receptor complexes, a mechanism that is difficult to achieve with antibodies optimized for direct competitive binding.

Crucially, the success of this work depended on rapid iteration between model generation, human structural inspection, and wet-lab feedback. Experimental results informed design decisions at multiple stages, including epitope definition, binder triage, and interpretation of apparent affinity gains. A seven-day binder validation pipeline enabled early rejection of unproductive binding modes and polyspecific binders, allowed testing of multiple hypotheses, and focused computational effort on experimentally viable solutions.

Beyond functional competition, this study also highlights a developability risk that is frequently underappreciated in the *de novo* miniprotein literature: molecular format can strongly amplify polyspecificity. Two designs exhibited low polyspecificity in both His-tagged and Fc-fusion contexts, whereas one design showed elevated off-target reactivity that increased markedly upon Fc fusion. Avidity-based increase in polyspecificity has been well documented for antibodies [32] Notably, Fc-mediated avidity-driven amplification was observed for CbAgo_b2, a previously published miniprotein with single-digit nanomolar affinity [6]. CbAgo_b2 has been reported to bind *Clostridium butyricum* Argonaute and inhibit its nuclease activity. In our assays, however, CbAgo_b2 exhibited pronounced polyspecificity, particularly when reformatted as a Fc fusion. While this does not negate its reported biochemical activity, it raises the possibility that some degree of functional inhibition may arise from nonspecific or avidity-enhanced interactions rather than from a uniquely precise binding mode. These results underscore that affinity alone is an insufficient screening endpoint for generative binder programs and motivate the incorporation of *in silico* prescreening strategies to detect off-target binding, including decoy-based structural confidence metrics (e.g., off-target iPTM) and physicochemical filters based on surface charge and hydrophobicity.

Collectively, these results establish epitope expansion as a generalizable strategy for targeting complex protein interfaces and underscore the importance of tightly coupled AI–human–experimental workflows in generative protein design. As the field moves toward increasingly ambitious targets characterized by multidomain architecture and high-affinity natural ligands, success will depend not only on improved models, but also on workflows that integrate rapid experimental validation and developability-aware screening early in the design process.

## Methods

### Latent-X1 computational design

*de novo* miniprotein binders were generated using the Latent-X1 Graphical User Interface Platform (https://platform.latentlabs.com/). Design targets were derived from experimentally determined IgE Fc structures (PDB: 8K7R), with truncations applied where necessary to satisfy model context-length constraints. Initial design campaigns targeted a nested epitope within the IgE C*_ε_*3 domain, followed by structure-guided epitope expansion toward the Fc*ε*RI-binding interface.

For each design batch, binder length ranged from length 100-150 residues, Latent-X1 generated candidate binder–target complexes that were scored using a composite metric integrating predicted TM-score (pTM), minimum inter-chain predicted aligned error (min iPAE), and complex RMSD following Chai-1 refolding. The composite score was defined as:

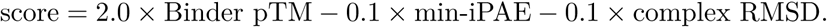

Top-ranking designs were visually inspected to confirm interface plausibility, absence of steric clashes with the full IgE Fc assembly, and compatibility with downstream experimental formats. Selected sequences were advanced to experimental validation. Visualization of complexes and structural analyses were performed using pymol.

### Rapid BLI binder validation

Initial experimental screening of designed miniproteins was performed using Onava’s high-throughput biolayer interferometry (BLI) workflow. Latent-X1–generated candidates, together with omalizumab and trastuzumab scFv controls, were codon optimized and formatted for cell-free expression using standardized DNA preparation protocols. Miniproteins were expressed using a cell-free protein synthesis system and processed in a parallelized, high-throughput format for BLI analysis. Binding measurements were acquired on a Gator Plus BLI instrument using a proprietary rapid-screening workflow optimized for early-stage binder validation. Human IgE Fc (Acro Biosystems) was used as the antigen at a concentration of 1 *µ*M, with an association phase of 200 s followed by a dissociation phase of 300 s.

### Dose-dependent kinetic validation of IgE binding

Dose-dependent kinetic samples were prepared similar to rapid binder analysis validation methods. Experiments were performed by immobilizing cell-free proteins on BLI biosensors at a constant capture level and challenging with serial dilutions of IgE starting with 1-2 uM concentration with a 2-fold serial dilution. Association (200 s) and dissociation phases (300 s) were fit using a 1:1 binding model where appropriate to extract apparent dissociation constants (*K_D_*), association rates (*k*_on_), and dissociation rates (*k*_off_). Experiments were performed in duplicates.

Purified proteins dose-response experiments were perform by immobilizing IgE Fc using anti-his probes or by immobilizing Fc-tagged proteins via protein A probes.

### Protein expression and purification

His-tagged miniproteins were expressed using a cell-free protein expression system and purified by Ni–NTA affinity chromatography. Fc-tagged constructs were expressed in Expi-CHO cells at 1 mL scale and purified using Protein A affinity chromatography, followed by size-exclusion chromatography (SEC) for high purity.

Protein purity and homogeneity were assessed by SEC. While most samples exhibited a dominant main peak, a few constructs displayed reduced main-peak fractions indicative of partial heterogeneity.

### Alanine-scanning mutagenesis

Alanine-scanning libraries were generated using the Onava platform mutagenesis and codon-optimization modules. Residues within 5 Å of the IgE-binding interface, as defined by Latent-X1 structural models, were individually mutated to alanine. All mutant constructs were codon optimized, expressed using the same cell-free protein synthesis workflow as the parental designs, and evaluated by BLI to quantify the impact of each substitution on IgE binding.

Binding effects were classified as deleterious (> 10*×* loss of affinity), neutral (within 10*×*), or affinity-enhancing (> 10*×* improvement) relative to the parental construct.

### BVP-ELISA polyspecificity profiling

Polyspecificity was assessed using a baculovirus particle (BVP) ELISA assay. His-tagged and Fc-tagged miniproteins were tested at 14 nM 70 nM, and 350 nM, Signals were normalized to highly polyreactive bococizumab highest dose. BVP-ELISA scores were calculated from triplicate measurements and used to assess format-dependent off-target reactivity.

### Polyspecificity BLI with VEGF isoforms

Polyspecificity BLI experiments were performed against the multidomain antigen VEGF165 and its truncated isoform VEGF110. The VEGF-binding miniprotein GDM_VEGFA_79 served as a positive control, while trastuzumab was included as a negative control. His-tagged miniproteins were immobilized to constant capture levels below 1.3 nm shift and subsequently challenged with 500 nM VEGF165 or VEGF110 (Acro Biosystems). Association and dissociation phases were conducted using the same timing parameters described for IgE-binding experiments.

### Competition BLI with Fc***ε***RI

Competition experiments were performed using a three-step tandem biolayer interferometry (BLI) assay. IgE Fc was first immobilized on biosensors, followed by association with Fc*ε*RI–Fc at saturating concentrations or buffer alone as a control during the first association step. In the second association step, Fc-tagged miniproteins were introduced at 2 *µ*M to assess binding to IgE in the presence or absence of pre-bound Fc*ε*RI.

## Acknowledgments

We thank Simon Kohl, Simon Mathis, and Daniella Pretorius for helpful discussions on deploying the Latent-X1 platform for protein binder design. We thank GenScript for protein production and characterization services. We also thank Danny Diaz, Engin Yapaci, Ravi Ramenani, Thanos Tsagkadouras, Michael Holden, Wang Cheng, Wayne Gu, Kirk Wiggan, Kadidiatou Diakite, Chetan Mishra, and Salvatore Candido for thoughtful discussions and feedback related to this manuscript.

