## Supplementary Data for "Iterative Epitope Expansion Enables Structure-Guided *de novo* Design of IgE-Binding Miniproteins Targeting the Fc*ε*RI Interface"

| Name | Expression system | Cal. M.W. (KDa) | Purification | Buffer | Concentration (mg/ml) | Purity by SDS-PAGE under R(%) | Size-Volume (ml) | Expression Volume (L) | Total (mg) |
|---|---|---|---|---|---|---|---|---|---|
| ONA-IgE-Rd1-2 | Cell-free system | 17.314 | Ni column | PBS, pH 7.2 | 1.31 | ≥85 | 0.8 | 0.005 | 1.04 |
| ONA-IgE-Rd1-3 | Cell-free system | 17.100 | Ni column | PBS, pH 7.2 | 1.37 | ≥55 | 0.8 | 0.005 | 1.09 |
| CbAgo_b2 | Cell-free system | 16.477 | Ni column | PBS, pH 7.2 | 1.22 | ≥70 | 0.8 | 0.005 | 0.976 |
| GDM_VEGFA_79 | Cell-free system | 14.113 | Ni column | PBS, pH 7.2 | 1.04 | ≥95 | 0.8 | 0.005 | 0.832 |
| Trastuzumab-scFv | Cell-free system | 27.088 | Ni column | PBS, pH 7.2 | 0.63 | ≥90 | 4.8 | 0.03 | 3.02 |
| Briakinumab-scFv | Cell-free system | 26.815 | Ni column | PBS, pH 7.2 | 0.38 | ≥80 | 4.8 | 0.03 | 1.82 |
| ONA-IgE-Rd2-1 | Cell-free system | 16.050 | Ni column | PBS, pH 7.2 | 0.90 | ≥90 | 0.8 | 0.005 | 0.72 |

**Supplementary Table 1. Summary of recombinant proteins produced using the cell-free expression system**. The table shows protein names, expression parameters, calculated molecular weight (M.W.), purification method, buffer composition, final protein concentration, purity assessed by SDS-PAGE under reducing conditions (R), final volume, expression volume, and total protein yield.

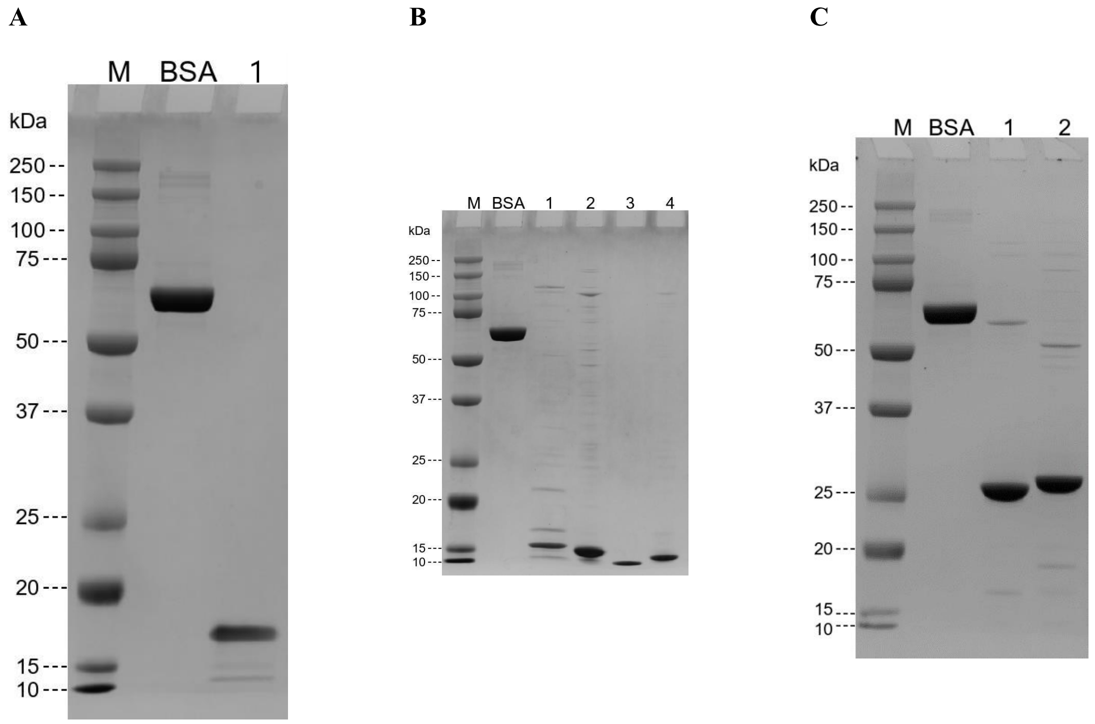


**Supplementary Fig. 1. SDS-PAGE analysis of purified proteins expressed in cell-free system under reducing conditions**. **(A)** Analysis of protein ONA-IgE-Rd1-2 (Lane 1). **(B)** Analysis of proteins ONA-IgE-Rd1-3, CbAgo_b2, GDM_VEGFA_79, and ONA-IgE-Rd2-1 (Lane 1-4). **(C)** Analysis of proteins Trastuzumab-scFv and Briakinumab-scFv (Land 1 and 2). For all panels: M indicates protein molecular weight marker; BSA lane contains 2.0 µg bovine serum albumin as a loading control. For each sample, 2 µg of purified protein or the maximal amount allowed was loaded per well. The position of molecular weight markers is indicated on the left side of each gel.

| Protein Name | Concentration (mg/ml) | Purity SDS-PAGE (%) | Purity SEC-HPLC (%) | Total(mg) |
|---|---|---|---|---|
| ONA-IgE-Rd1-2-Fc | 7.28 | 95 | 93 | 8.01 |
| ONA-IgE-Rd1-3-Fc | 4.04 | 95 | 60 | 4.44 |
| CbAgo_b2-Fc | 2.68 | 95 | 73 | 2.95 |
| GDM_VEGFA_79-Fc | 0.39 | 80 | 66 | 0.43 |
| Trastuzumab | 5.99 | 90 | 94 | 7.79 |
| Briakinumab | 4.06 | 80 | 95 | 5.28 |
| ONA-IgE-Rd2-1-Fc | 0.83 | 95 | 65 | 0.91 |

**Supplementary Table 2. Summary of recombinant proteins produced using the CHO (Chinese Hamster Ovary) cell expression system.** The table presents protein names, final concentration, purity assessed by SDS-PAGE (sodium dodecyl sulfate polyacrylamide gel electrophoresis), purity assessed by SEC-HPLC (size exclusion chromatography-high performance liquid chromatography), and total protein yield. N/A indicates that purity assessment was not applicable or not performed for that particular sample.

### A. ONA-IgE-Rd1-2-Fc

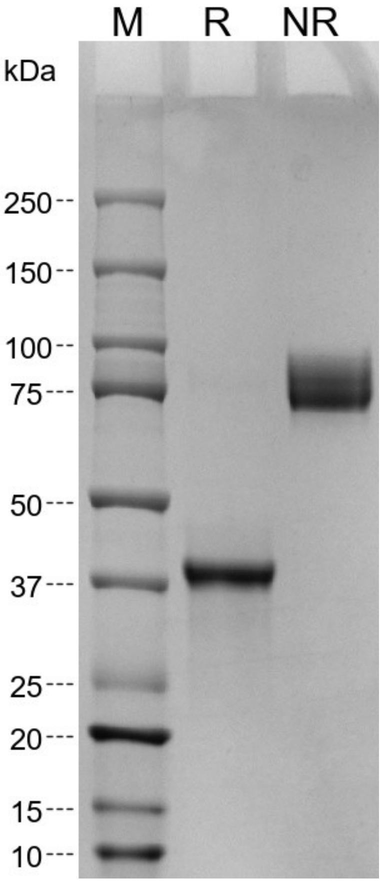


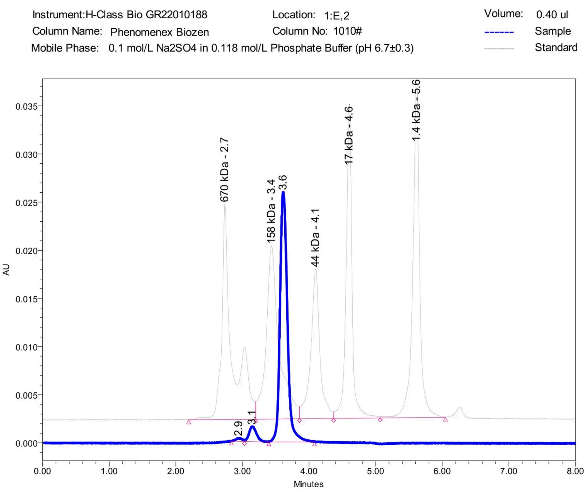


| | RT | Height | Width (sec) | Area | AreaPercentage |
|---|---|---|---|---|---|
| 1 | 2.9 | 342.1 | 11.9 | 2286.9 | 1 |
| 2 | 3.1 | 1548.7 | 22.0 | 12393.1 | 6 |
| 3 | 3.6 | 25970.2 | 41.1 | 200631.7 | 93 |

**B. ONA-IgE-Rd1-3-Fc**

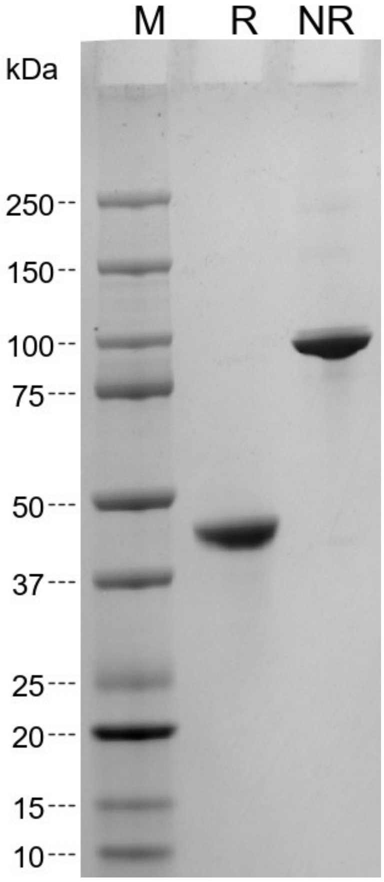


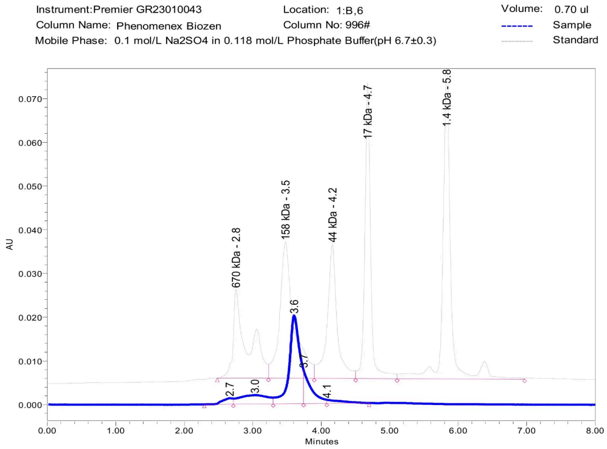


| | RT | Height | Width (sec) | Area | AreaPercentage |
|---|---|---|---|---|---|
| 1 | 2.7 | 1365.1 | 25.6 | 13959.9 | 4 |
| 2 | 3.0 | 2008.3 | 34.9 | 59206.1 | 16 |
| 3 | 3.6 | 20168.0 | 26.6 | 214737.2 | 60 |
| 4 | 3.7 | 7730.3 | 20.4 | 58878.0 | 16 |
| 5 | 4.1 | 811.0 | 37.1 | 12293.8 | 3 |

**C. CbAgo_b2-Fc**

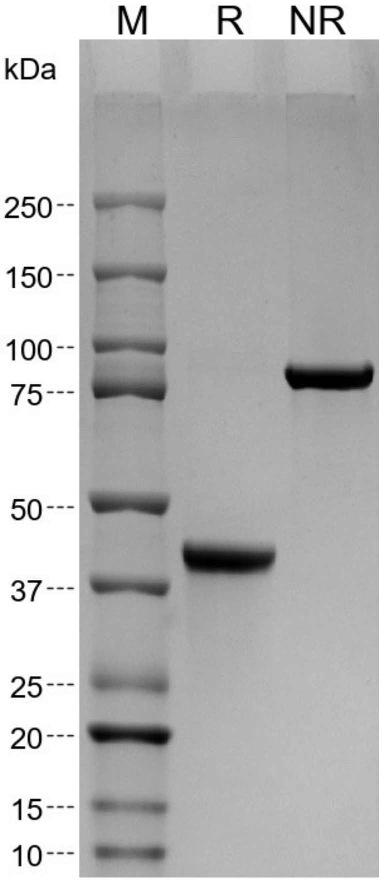


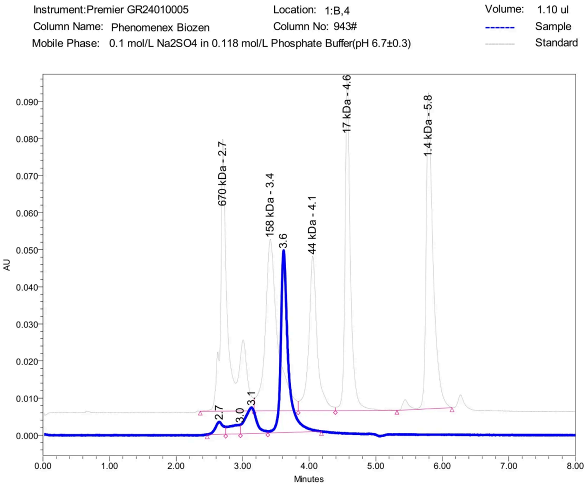


| | RT | Height | Width (sec) | Area | AreaPercentage |
|---|---|---|---|---|---|
| 1 | 2.7 | 3352.1 | 16.6 | 28416.1 | 5 |
| 2 | 3.0 | 2445.2 | 13.4 | 29310.6 | 6 |
| 3 | 3.1 | 7012.2 | 24.6 | 83224.8 | 16 |
| 4 | 3.6 | 49121.0 | 48.5 | 380936.4 | 73 |

**D. GDM_VEGFA_79-Fc**

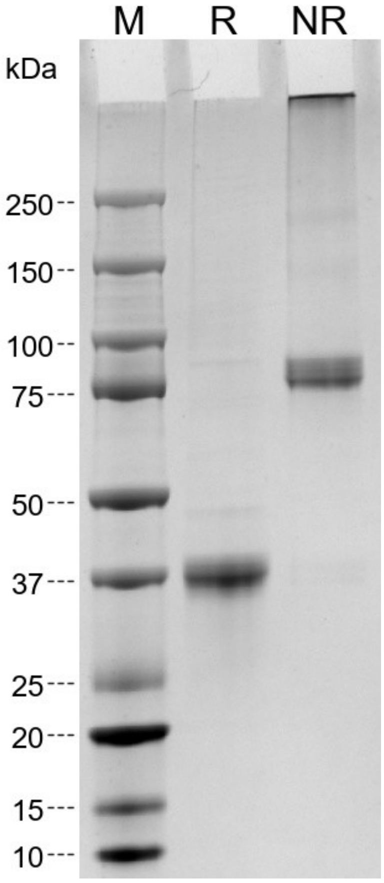


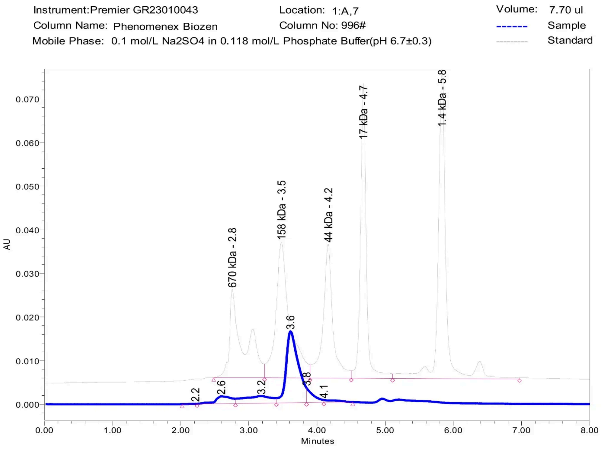


| | RT | Height | Width (sec) | Area | AreaPercentage |
|---|---|---|---|---|---|
| 1 | 2.2 | 81.5 | 13.1 | 608.3 | 0 |
| 2 | 2.6 | 1677.2 | 34.0 | 28673.2 | 9 |
| 3 | 3.2 | 1642.6 | 35.9 | 47718.6 | 15 |
| 4 | 3.6 | 16376.3 | 26.5 | 205935.2 | 66 |
| 5 | 3.8 | 3379.4 | 15.3 | 23700.3 | 8 |
| 6 | 4.1 | 548.4 | 25.6 | 7202.9 | 2 |

### E. Trastuzumab

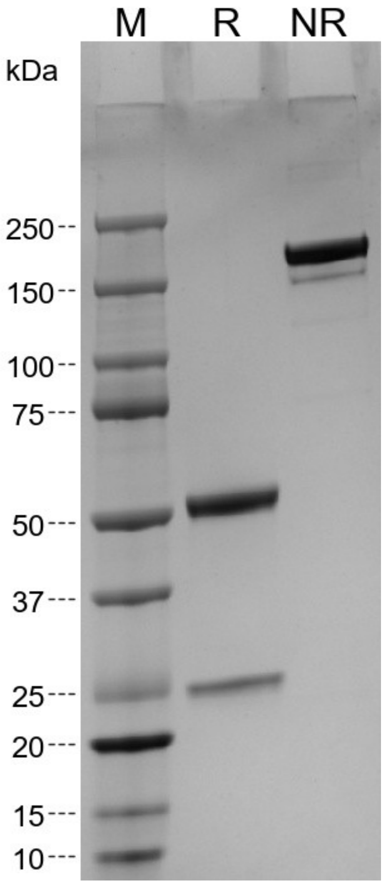


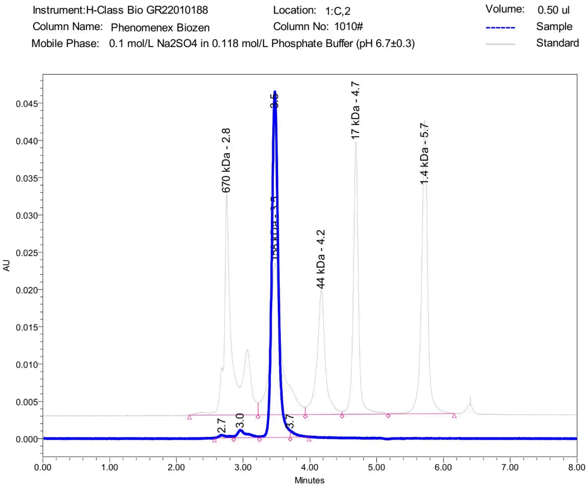


| | RT | Height | Width (sec) | Area | AreaPercentage |
|---|---|---|---|---|---|
| 1 | 2.7 | 400.0 | 17.3 | 3760.6 | 1 |
| 2 | 3.0 | 1036.9 | 23.3 | 10762.1 | 3 |
| 3 | 3.5 | 46382.8 | 27.6 | 309982.2 | 94 |
| 4 | 3.7 | 872.3 | 17.0 | 5067.6 | 2 |

### F. Briakinumab

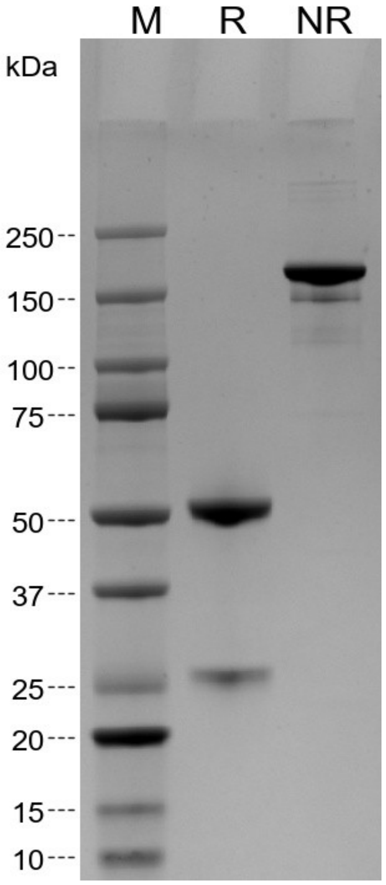


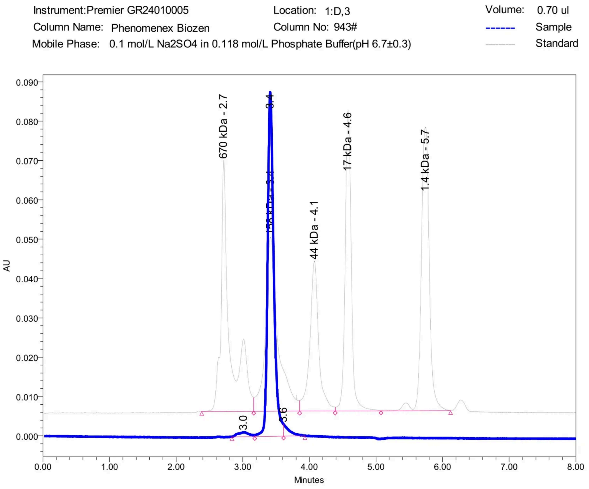


| | RT | Height | Width (sec) | Area | AreaPercentage |
|---|---|---|---|---|---|
| 1 | 3.0 | 1147.9 | 20.6 | 13709.5 | 2 |
| 2 | 3.4 | 87472.0 | 25.9 | 558211.5 | 95 |
| 3 | 3.6 | 3005.4 | 19.2 | 17624.8 | 3 |

### G. ONA-IgE-Rd2-1-Fc

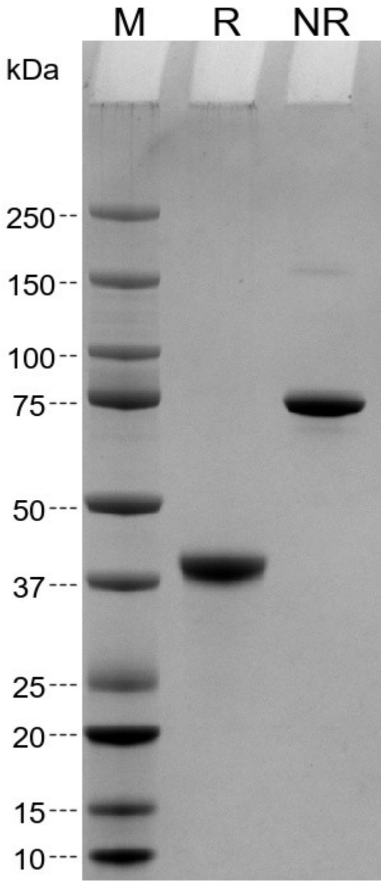


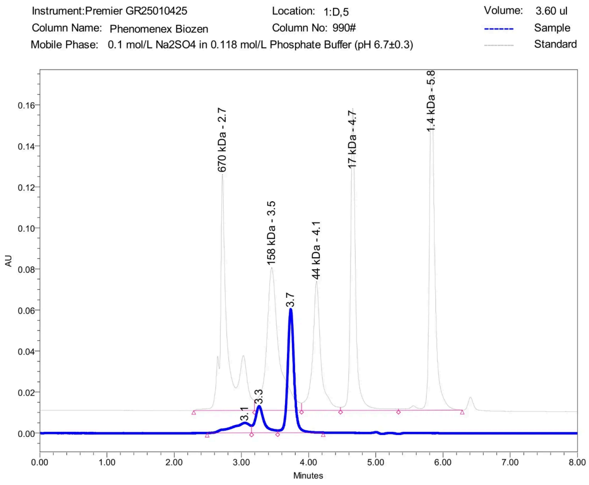


| | RT | Height | Width (sec) | Area | AreaPercentage |
|---|---|---|---|---|---|
| 1 | 3.1 | 4868.5 | 39.7 | 93341.0 | 15 |
| 2 | 3.3 | 12995.0 | 23.3 | 118549.7 | 20 |
| 3 | 3.7 | 60157.3 | 40.6 | 393987.3 | 65 |

**Supplementary Fig. 2. Quality control analysis of CHO cell-expressed proteins.** For each protein **(A-G)**, SDS-PAGE analysis (left panel) and SEC-HPLC chromatogram (right panel) are shown. **SDS-PAGE panels:** M indicates protein molecular weight marker; R indicates sample run under reducing conditions; NR indicates sample run under non-reducing conditions. **SEC-HPLC panels:** X-axis shows retention time (minutes); Y-axis shows absorbance unit (AU). The major peak represents the monomeric protein, while smaller peaks or shoulders indicate aggregates or degradation products. The percentage of monomeric protein is calculated from the area under the main peak relative to the total peak area.
